# Dynamic interplay of protrusive microtubule and contractile actomyosin forces drives tissue extension

**DOI:** 10.1101/2022.06.21.496930

**Authors:** Amrita Singh, Sameedha Thale, Tobias Leibner, Andrea Ricker, Harald Nüsse, Jürgen Klingauf, Mario Ohlberger, Maja Matis

## Abstract

In order to shape a tissue, cell-based mechanical forces have to be integrated into global force patterns. Over the last decades, the importance of actomyosin contractile arrays, which are the key constituents of various morphogenetic processes, has been established for many tissues. Intriguingly, recent studies demonstrate that the microtubule cytoskeleton mediates folding and elongation of the epithelial sheet during *Drosophila* morphogenesis, placing microtubule mechanics *en par* with actin-based processes. While these studies establish the importance of both cytoskeletal systems during cell and tissue rearrangements, a mechanistic explanation of their functional hierarchy is currently missing. Here, we dissect the individual roles of these two key generators of mechanical forces during epithelium elongation. We demonstrate that microtubules dictate cell shape changes and actomyosin refines them. Combining experimental and numerical approaches, we find that altering the microtubule and actomyosin functions results in predictable changes in tissue shape. We further show that planar polarized microtubule patterning is independent of cell geometry and actomyosin-based mechanics. These results support a hierarchical mechanism, whereby microtubule-based forces in some epithelial systems prime actomyosin-generated forces.

## Introduction

Tissue morphogenesis results from a finely tuned spatial and temporal integration of various cellular behaviors, including changes in cell shape and size, cell migration, division, and cell intercalation (Gilmour et al., 2017). These distinct behaviors are driven by tissue intrinsic and extrinsic mechanisms, which jointly coordinate mechanical forces exerted on cells during tissue patterning (Collinet and Lecuit, 2021, Heisenberg and Bellaiche, 2013, Pinheiro and Bellaiche, 2018). Within individual cells, actomyosin filaments together with microtubules and intermediate filaments form the composite cytoskeleton that controls cell mechanics during tissue remodeling. While studies have already established the importance of actin-based mechanical forces coupled via intercellular adherens junctions (Clarke and Martin, 2021), relatively little is known about the contribution of other cytoskeletal components to cell shape changes and cell mechanics during morphogenesis. Microtubules were initially considered to participate only in a supporting role in cell mechanics, contributing, for instance, to the stabilization of the actomyosin or trafficking of adhesion molecules (Bouchet and Akhmanova, 2017, Etienne-Manneville, 2013, Stehbens and Wittmann, 2012). However, recent work has challenged this view, demonstrating that microtubules are capable of generating protrusive forces that are crucial for key morphogenetic processes, including tissue bending and tissue extension (Singh et al., 2018, Takeda et al., 2018, Matis, 2020). Yet, while these studies demonstrate that actomyosin and microtubule-based forces are equally important, it remains unclear how these force-generating systems interact. For instance, how are different forces directed in space and time, and what is the hierarchy between them?

Here, we used the fruit fly *Drosophila melanogaster* wing development, a versatile model system for *in vivo* studies of force patterning, to probe the interplay between actin and microtubule dynamics. During pupal wing morphogenesis, the epithelium initially displays substantial cell shape changes (14-18 hours After Puparium Formation; hAPF), followed by cell rearrangements (22-28 hAPF) (Classen et al., 2005, Aigouy et al., 2010, Bardet et al., 2013).

Initial cell elongation during phase I (14-18 hAPF) relies on the coordinated generation and integration of local microtubule-based forces into a global tissue force pattern along the proximal-distal axis (Singh et al., 2018). The proximal-distal alignment of microtubules results from the planar polarization of the tissue through the Fat planar cell polarity (Fat-PCP) signaling pathway (Harumoto et al., 2010, Matis et al., 2014, Olofsson et al., 2014). Fat-PCP consists of the atypical cadherins, Fat (Ft) and Dachsous (Ds), and the Golgi resident protein Four-jointed (Fj), a transmembrane kinase (Zeidler et al., 2000, Villano and Katz, 1995, Yang et al., 2002, Ma et al., 2003). Ft and Ds interact across adherens junctions, forming heterodimers across adjacent cells that Fj modulates. As Fj and Ds expression both display gradients along the proximal-distal axis, Ft–Ds heterodimers accumulate in a polarized fashion along the global axis. The net result is the conversion of tissue-wide transcriptional gradients of Fj and Ds into functional polarization of Ft-Ds at the cellular level, thereby providing a spatial cue to orient diverse developmental processes. Consistently, Ft-PCP was shown to orient cell divisions, tissue growth, cell rearrangements and cell migration in *Drosophila*, zebrafish and mammals (Zakaria et al., 2014, Mao et al., 2016, Saburi et al., 2008, Baena-Lopez et al., 2005, Aigouy et al., 2010, Mao et al., 2011, Chen et al., 2018, Durst et al., 2015, Li-Villarreal et al., 2015). Unlike the initial steps described above, wing tissue remodeling during phase II (22-28 hAPF) depends on extrinsic tensile forces generated by wing hinge contraction that starts at 18 hAPF (Etournay et al., 2015, Ray et al., 2015, Sugimura and Ishihara, 2013, Aigouy et al., 2010). Here, an apical extracellular matrix protein Dumpy anchors the wing epithelium to the adjacent cuticle (Ray et al., 2015, Etournay et al., 2015). The attachment of the wing tissue to the cuticle leads to resistance to pulling forces generated by hinge constriction, resulting in tension along the proximal-distal axis. Subsequently, the buildup tension drives wing remodeling, including cell shape changes, oriented cell divisions and rearrangements along the same axis (Aigouy et al., 2010, Sugimura and Ishihara, 2013).

In this study, we focus on the interplay of actin and microtubule-based forces during phase I (Supp Figure 1A). While previous studies establish the presence of both systems during early pupal wing morphogenesis (Singh et al., 2018, Sugimura and Ishihara, 2013), the exact contribution of microtubule and MyosinII-generated forces to cell and wing shape remains an open and intriguing question. Our work uncovers that the balance of actomyosin and microtubule-generated forces control cell lengthening during epithelium development. The proposed mechanism controlling local mechanics, which in turn coordinates cell behaviors, is likely relevant to many other tissues that are planar polarized by the Ft-PCP signaling pathway.

## Results

### MyosinII anisotropy leads to junctional tension pattern

As previously demonstrated (Singh et al., 2018), MyosinII localizes during phase I (14-18 hAPF) to the apical cell junctions, where it is planarly polarized along the longer (proximal-distal) axis of the elongating cells, forming anisotropic cables across the entire wing tissue (Figure 1A). Intriguingly, ablated junctions aligned along the proximal-distal axis retract stronger upon incision than junctions aligned along the anterior-posterior axis (Figure 1B and Supp Figure 1B,C), arguing for an anisotropic distribution of tension at adherens junctions.

**Figure 1.**
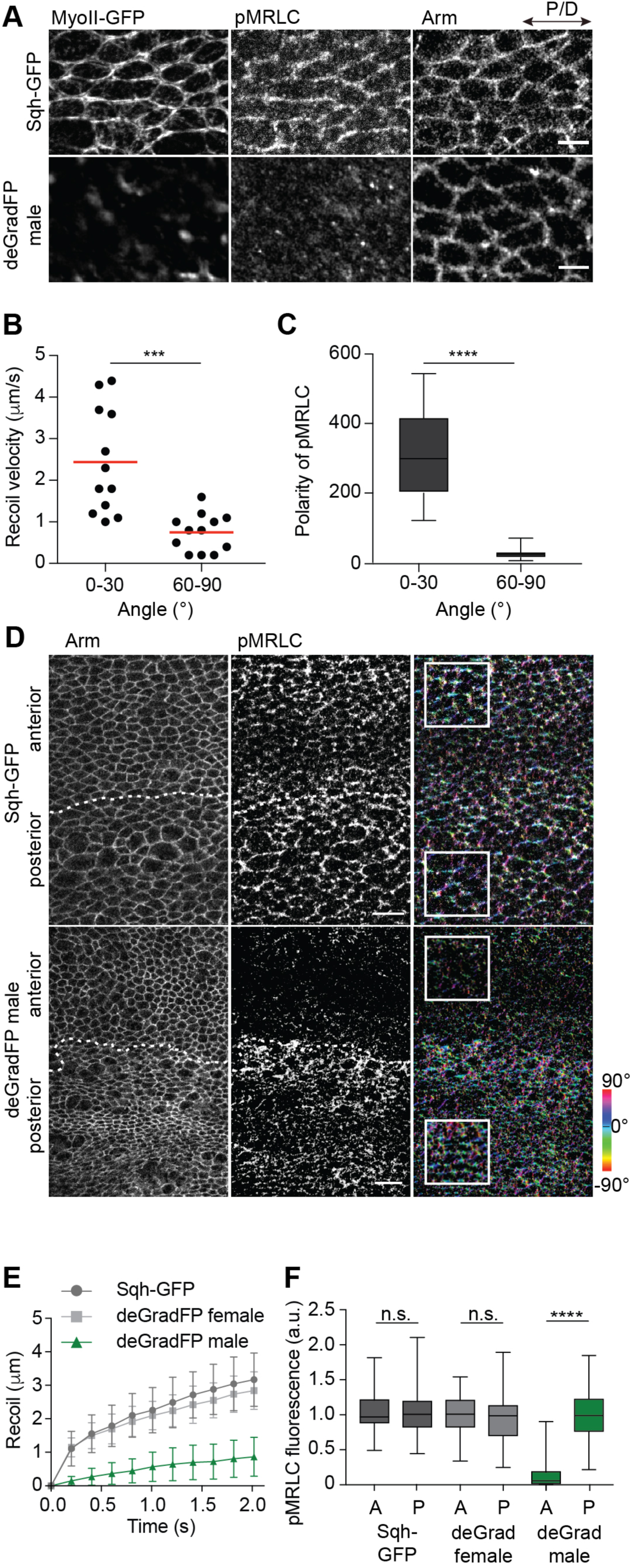
MyosinII is polarized during early pupal wing development. (**A**) Representative images showing anterior compartment of Sqh-GFP male (top) and deGradFP male (lower). To get a developmental stage equivalent to 18 hAPF at 25°C, deGradFP wings were grown sequentially at 18°C and 29°C using the tub-Gal80^ts^ system (see table S1). Wings are expressing Sqh-GFP and are stained for pMRLC and Arm. In wild-type flies (Sqh-GFP male), MyosinII and phosphomyosin show polarized distribution at junctions oriented along the proximal-distal axis. In deGradFP-expressing flies, Sqh-GFP and the phosphomyosin signal are lost from junctions. The remaining Sqh-GFP signal does not colocalizes with the phosphomyosin staining. Images shown in (A) are representative of n=5 wings and N=3 independent experiments. Scale bars, 5 µm. (**B**) Quantification of recoil velocity upon laser ablation of P/D (0-30°) and A/P (30-60°) interfaces in 18 hAPF wild-type flies (two-tailed t-test with Welch’s correction, *** p = 0.0007, n=12 junctions each and N=5 pupae). (**C**) Quantification of pMRLC polarity (Mann-Whitney test, **** p < 0.0001, n=450 cells and 4 pupae). (**D**) Images showing anterior-posterior border in control flies (Sqh-GFP male, top) and Sqh-GFP knockdown flies (deGradFP male, bottom) stained for Arm and pMRLC. In control flies, phosphomyosin localizes to the junctions in the anterior and posterior compartments. In the flies expressing deGradFP under *ci-Gal4* control, the phosphomyosin signal in the anterior compartment is lost compared to the wing’s posterior site. The polarity of the pMRLC signal is color-coded using OrientationJ. Scale bar, 10 µm. (**E**) Quantification of displacement upon laser ablation for interfaces along the proximal-distal axis in Sqh-GFP and female deGradFP control flies (gray, light gray) and Sqh-GFP knockdown flies (deGradFP male, green). n =10 junctions and 4 pupae per genotype. Values on the graph show mean displacements. Error bars show sd. (**F**) Quantifications of mean intensities of pMRLC in anterior and posterior compartments for Sqh-GFP and female deGradFP control flies (gray, light gray) and Sqh-GFP knockdown flies (deGradFP male, green). (Kruskal-Wallis tests, from left to right: n.s. p > 0.9999, n.s. p > 0.9999, **** p < 0.0001, number of junctions for Sqh-GFP male (A)=90, (P)=77, deGradeFP female (A)=76, (P)=74 and for deGradFP male (A)=100, (P)=102 and 3-5 pupae per genotype). Boxes in the plot extend from the 25^th^ to 75^th^ percentiles, with a line at the median. Whiskers show min and max values.

**Table 1.** A table summarizing the dual temperature regimes deployed for the analysis of different parameters using the deGradFP system.

| Temperature regimes<br>(at 18°C+29°C) | Equivalent stage<br>(at 25°C) | Assay(s) done using the regime | Explanation of the regime (in brief) |
| --- | --- | --- | --- |
| 22+6.5 hrs | 18 hAPF | • Standardisation for 18 hAPF pupal wing.<br>• Tension analysis (laser ablation).<br>• Cell shape and area analysis.<br>• MT alignment assay. | 0 hAPF pre-pupae collected and grown at 18°C for 22 hrs followed by incubation at 29°C for 6.5 hrs. |
| 22+14 hrs | 24 hAPF | • PCP organisation and cell area analysis<br>• Vein area analysis | 0 hAPF pre-pupae collected and grown at 18°C for 22 hrs followed by incubation at 29°C for 14 hrs. |
| 22+20 hrs | 30 hAPF | • Pupal wing hair polarity | 0 hAPF pre-pupae collected and grown at 18°C for 22 hrs followed by incubation at 29°C for 20 hrs. |
| 28+3 hrs | N/A | • Adult wing hair polarity.<br>• Adult wing morphology. | 0 hAPF pre-pupae collected and grown at 18°C for 22 hrs followed by incubation at 29°C for 3 hrs. |
| 0+24 hrs | N/A | • Cell division defects. | 0 hAPF pre-pupae collected and grown at 29°C for 24 hrs. |

To further explore this observation, we visualized the active phosphorylated form of myosin light chain (pMRLC), a bona fide marker for actomyosin contractility (Ikebe and Hartshorne, 1985). Notably, in 18 hAPF wings, pMRLC is enriched at the apical region and associated with cell junctions (Figure 1A). Consistently, quantification of pMRLC polarity demonstrated significant enrichment of pMRLC at proximal-distal oriented junctions that recoiled faster upon ablation (Figure 1C). Although MyosinII anisotropy positively correlates with the junctional tension pattern, it does not account for the possibility that unconventional myosin motor proteins may also contribute to junctional contractility (Bosveld et al., 2012, Lin et al., 2007). To account for this possibility, we selectively disrupted MyosinII function and assessed its effects on cell shape changes during early developmental stages. This was accomplished by taking advantage of a degron-based protein knockdown system (deGradFP) (Caussinus et al., 2011), which was efficiently used to deplete the GFP-tagged MyosinII light chain Sqh (Sqh-GFP) at the different developmental stage (Caussinus et al., 2011, Pasakarnis et al., 2016). Since expression of deGradFP resulted in early embryonic lethality of males, we used the tub-Gal80^ts^ system that allows a temporal regulation of gene expression. To ensure that the final developmental stage was equivalent to 18 hAPF at 25°C, we first standardized the growth protocol in the presence of a temperature shift (i.e., 18°C to 29°C, see Table 1, Supplemental information). Next, the selected temperature regime was validated by quantifying cell elongation and apical cell area (Supp Figure 2A-D). Expression of the deGradFP system disrupted actomyosin cables in MyosinII depleted cells (Figure 1A,D). To test for changes in tension at adherens junctions along the proximal-distal axis, we next performed laser ablation experiments (Supp Figure 2E). Compared to controls (Sqh-GFP male and deGradFP female), deGradFP males displayed a striking drop in recoil velocity (Figure 1E), indicative of reduced tension along the MyosinII enriched junctions. Consistently, deGradFP mediated knockdown showed a strong reduction of overall MyosinII phosphorylation (Figure 1D,F). Notably, deGradFP-mediated knockdown of Sqh-GFP presented phenotypes reminiscent of MyosinII loss-of-function mutants in *Drosophila* (Supp Figure 3). Finally, deGradFP females, which carry a wild-type copy of MyosinII, show no defects observed in deGradFP males providing the evidence that the system does not cause any MyosinII-unspecific phenotypes (Supp Figure 4). Collectively, these results establish that MyosinII promotes the formation of the polarized junctional tension along the main tissue axis and excludes the possible contribution of unconventional myosin motors to patterned junctional tension.

### The interplay between microtubule and actomyosin regulates cell shape

Initial cell elongation during pupal wing development takes place between 14 to 18 hAPF (Aigouy et al., 2010, Singh et al., 2018). To test whether polarized MyosinII also participates in this process, we examined cell shape changes upon MyosinII inhibition (Figure 2A). Strikingly, quantification of cell length showed that cells depleted of MyosinII were significantly longer than control cells (Figure 2B, green). Together with the observed MyosinII polarization along proximal-distal junctions (Figure 1A), these findings suggest that MyosinII mediates contractile stress to limit the cell length. In addition, it posits that Myosin-independent forces are needed for cell elongation.

**Figure 2.**
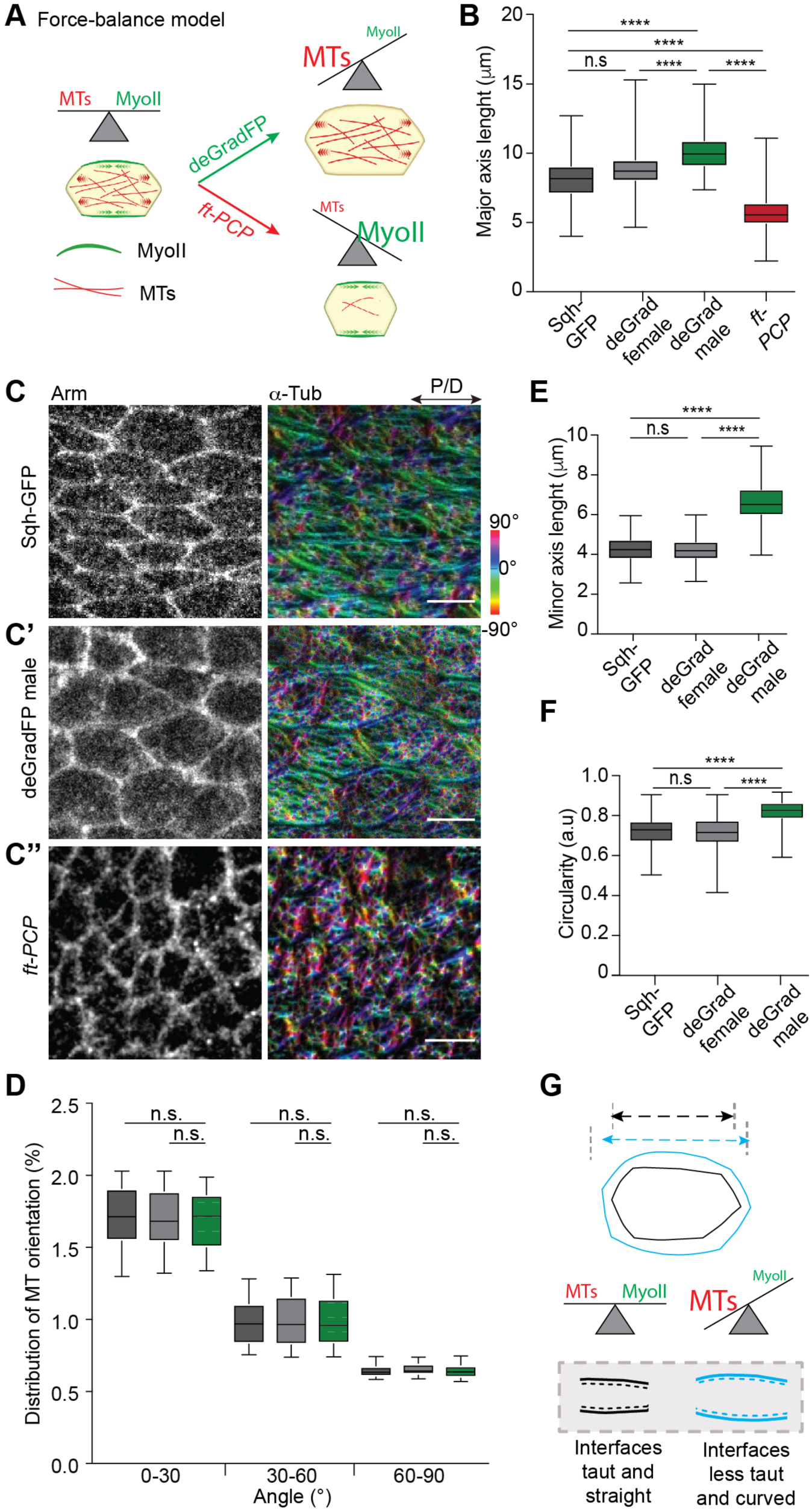
Polarized alignment of apical microtubules is not disrupted by the loss of MyosinII activity. (**A**) Force-balance model depicting the effect on cell shape/elongation upon disruption of MyosinII and microtubule forces. (**B**) Quantifications of cell length for different genotypes (Kruskal-Wallis test, n.s. p > 0.9999, **** p < 0.0001, **** p < 0.0001, p < 0.0001, n(control/ Sqh-GFP male)=497 cells, n(deGradFP female)=250, n(deGradFP male)=250 and n(*ft-PCP*)=261 cells and 3-5 pupae per genotype). (**C-C’’**) Images of 18 hAPF wings of Sqh-GFP male (C), deGradFP male (C’) and *ft-PCP* (*ft^l(2) fd^/ft^GRV^;tub-Gal4/UAS-Ft*Δ*ECD*Δ*N-1,* C’’) stained for Arm and α-Tub to visualize microtubules. Orientation of microtubules is color-coded using OrientationJ. Images shown are representative of 4 wings and 3 independent experiments. Scale bars, (C-C’’) 5 µm. (**D**) Quantification of microtubule alignment along proximal-distal axis for control flies (Sqh-GFP male and deGradFP female; gray, light gray) and deGradFP male (Kruskal-Wallis tests: n.s. p > 0.9999, n=80-100 cells and 3 pupae per genotype). (**E**) Quantifications of circularity for different genotypes. (Kruskal-Wallis tests from left to right: **** p < 0.0001, n.s. p > 0.9999, **** p < 0.0001, n(control)=250 cells, n(deGradFP female)=250 cells, n(deGradFP male)=250 cells and 3-5 pupae per genotype). Boxes in all plots extend from the 25^th^ to 75^th^ percentiles, with a line at the median. Whiskers show min and max values. (**F**) Quantifications of cell minor axis for different genotypes (Kruskal-Wallis tests from left to right: **** p < 0.0001, n.s. p > 0.9999, **** p < 0.0001, n(control)=250 cells, n(deGradFP female)=250 cells, n(deGradFP male)=250 cells and 3-5 pupae per genotype). (**G**) Cartoon showing the effect of disruption of MyosinII and microtubule activity on cell shape. The cell shape changes result from a general release of cellular pre-stress upon loss of MyosinII contractility, as suggested by an increase of length and width of the cell. Thus, indicating there is no direct relation between MyosinII polarized organization and cell elongation along the proximal-distal axis. However, MyosinII refines the overall cell shapes by (i) regulating the aspect ratio of the cells and (ii) by keeping the interfaces along proximal-distal axis taut and straight (indicated by black broken lines) as shown through the region of cell interfaces marked within the gray box (G, bottom).

As previously described, wing cell elongation entails microtubule alignment along the proximal-distal axis (Singh et al., 2018), whereby microtubule patterning is regulated through the Fat-PCP signaling pathway (Harumoto et al., 2010, Matis et al., 2014, Olofsson et al., 2014). We thus hypothesized that in MyosinII depleted cells, where cortical tension is reduced, microtubule-generated protrusive forces may drive cell elongation. Should this indeed be the case, polarized microtubule patterning should also occur in deGradFP flies lacking actomyosin contractility. Strikingly, microtubule alignment along the proximal-distal axis was comparable for deGradFP and control flies (Figure 2C-C’). Quantification of the angular distribution of microtubules with respect to the proximal-distal axis revealed that in deGradFP flies, 84% of all microtubules aligned within 0-30° from the principal proximal-distal axis, which is analogous to control cells (85%) (Figure 2D). These data suggest that patterned microtubule cytoskeleton directs cell shape changes. To further strengthen the validity of this hypothesis, we probed whether misalignment of microtubules results in shorter cells (Figure 2A). Indeed, cells in *ft* mutant animals rescued for the Hippo pathway (hereafter called *ft-PCP* mutant), in which microtubules are misoriented (Figure 2C’’), display a smaller cell elongation index (EI, defined by the ratio of the length of the longest cell axis to the shortest cell axis) (Singh et al., 2018). Considering that an increase in cell width and a decrease in cell length both lower the EI, we next measured the absolute cell length in the *ft-PCP* mutant. Consistent with the role of aligned microtubules for cell elongation, we find that cells indeed become shorter in the *ft-PCP* mutant background (Figure 2B, red). Finally, as disrupted cell polarity in the *ft-PCP* mutant may lead to cell shape changes independently of microtubule-generating forces, we took advantage of Patronin to specifically perturb microtubule organization. Members of the calmodulin-regulated spectrin-associated protein (CAMSAP) family in vertebrates and Patronin in invertebrates play an essential role in organizing microtubule cytoskeleton in several differentiated cells (Noordstra et al., 2016, Takeda et al., 2018, Toya et al., 2016, Panzade and Matis, 2021, Nashchekin et al., 2016). Consistently, Patronin knockdown in wing cells resulted in a dramatic change in the organization of apical non-centrosomal microtubules (Figure 3A,B). Importantly, Patronin depleted cells were also shorter (Figure 3C). Collectively, these independent experimental approaches consistently demonstrate that patterning of non-centrosomal microtubules mediates cell elongation.

**Figure 3.**
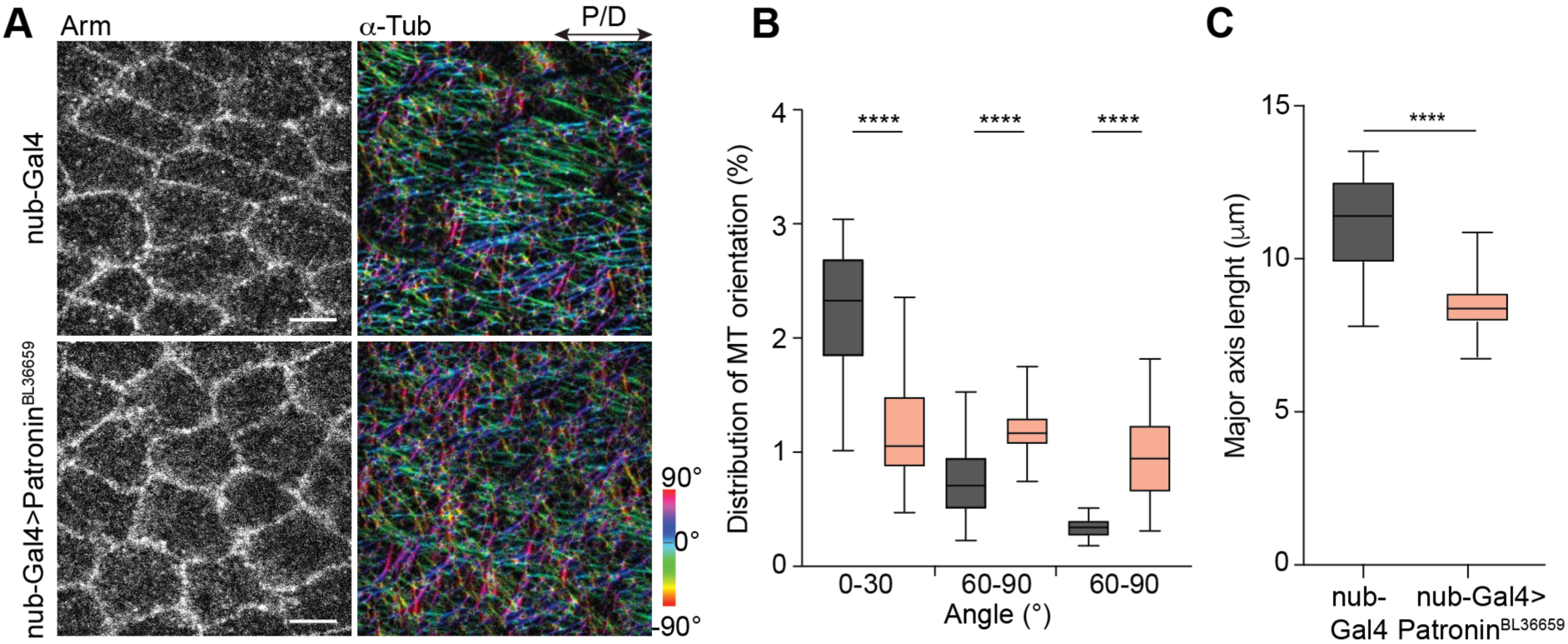
Depletion of Patronin perturbs microtubule organization and leads to shorter cells and tissue. (**A**) Images of 18 hAPF control (*nub-Gal4*; top) and Patronin depleted wings (*nub-Gal4>Patronin^RNAi^*; bottom) stained for Arm and α-Tub to visualize microtubules. The orientation of microtubules is color-coded using OrientationJ. The images shown are representative of 4 wings and 3 independent experiments. Scale bars, 5 µm. (**B**) Quantification of microtubule alignment along proximal-distal axis for control and Patronin depleted cells (Kruskal-Wallis tests, from left to right: **** p < 0.0001, **** p < 0.0001, **** p < 0.0001, n=80-100 cells and 3 pupae). (**C**) Quantifications of cell length. (Mann-Whitney test, **** p < 0.0001, n(control)=40 cells, n(*nub-Gal4>Patronin^RNAi^*)=38 cells and 3 pupae per genotype). Boxes in the plot extend from the 25^th^ to 75^th^ percentiles, with a line at the median. Whiskers show min and max values.

Having established that MyosinII is dispensable for initial cell elongation, we took a closer look at its role in other aspects of cell shape changes. MyosinII appeared to be required to refine the final cell shape (Figure 2C’). This was evident from the change in curvature of cellular interfaces in deGradFP males, where MyosinII contractility was disrupted, and cortical tension was released. In addition, cells in MyosinII-depleted wings expanded and widened along the anterior-posterior axis (Figure 2E) leading to cell rounding (Figure 2F). Hence, our results demonstrate that polarized MyosinII contractility is not required for initial polarized cell elongation along the proximal-distal axis but is needed for the subsequent refinement of cell shape (Figure 2G).

### A model of force balance predicts polarized microtubule-generated protrusive forces as an essential driver of cell elongation

We aimed to formalize these findings in a force-balance model, whereby microtubule-generated protrusive forces counterbalanced by contractile actomyosin forces drive cell elongation (Figure 2A). Various continuum and agent/vertex-based models have been devised on the cell level and validated in different context (Alt et al., 2017). Further, attempts have been undertaken to rigorously derive macroscopic continuum models on the tissue level, starting from individual cell-based models (Penta et al., 2014). In most of these settings, however, the effects of the microtubule or cytoskeleton reorganization are not taken into account. We therefore developed and implemented a continuum model, which is adapted to our setting. Since in our case the cells are mechanically autonomous (Singh et al., 2018), it suffices to model a single cell. To average out microscopic details, while maintaining the structural properties, the model regards the cell from a mesoscopic point of view (Figure 4A). The microtubule cytoskeleton is modeled as a viscoelastic gel formed by polar filaments that can actively exert forces (active polar gel) (Fan and Li, 2015). The actomyosin cortex is modelled through the effective tension of the cell surface. For full details on the mathematical model, its efficient numerical discretization and implementation see (Leibner et al., 2021) and STAR methods.

**Figure 4.**
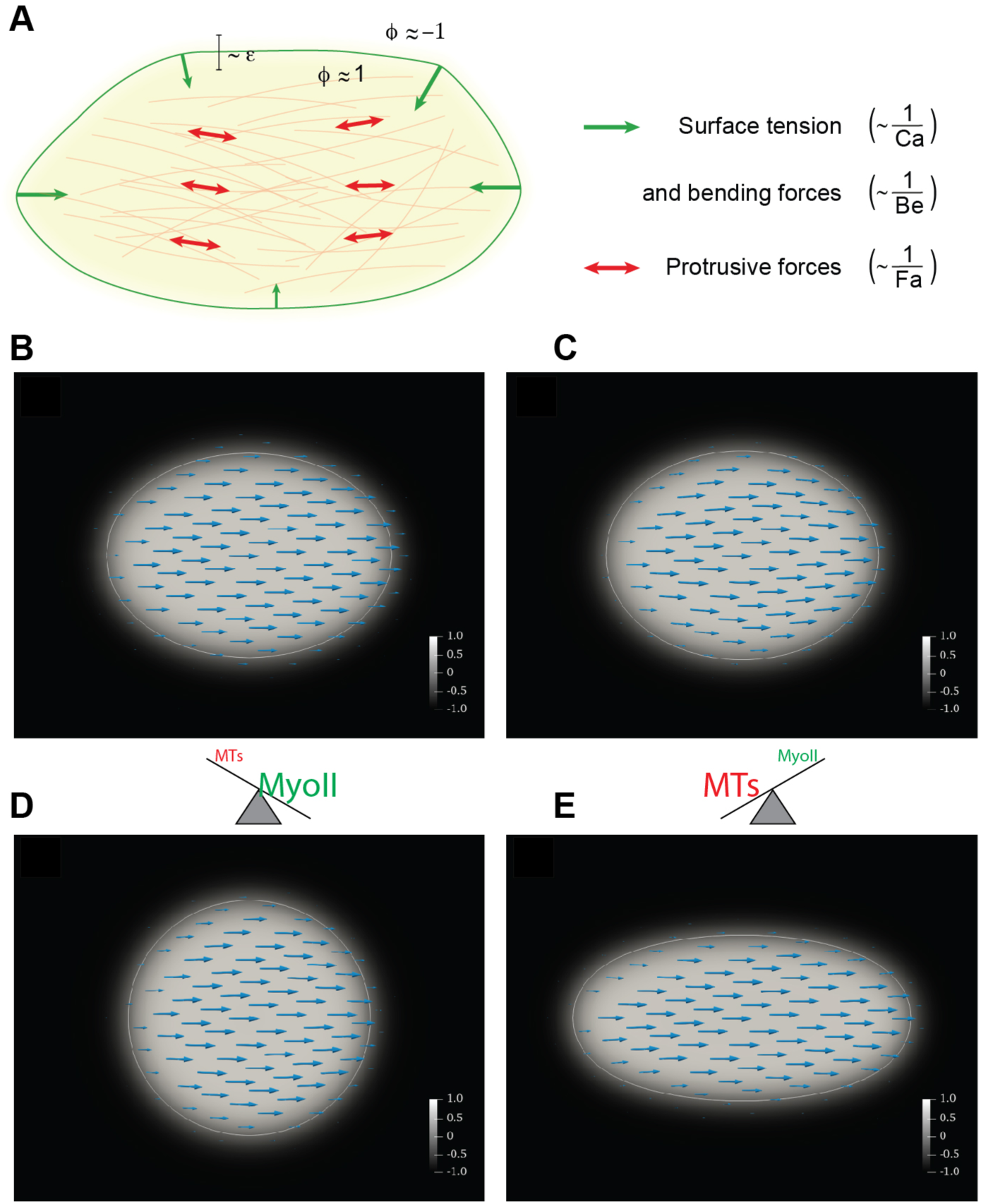
Computational verification of the force-balance hypothesis. (**A**) Schematic description of the computational model. The cell is modeled by the phase field variable ϕ that takes on the value 1 inside the cell (beige area) and -1 outside with a smooth transition in the interface region whose thickness is proportional to the parameter ɛ. The cell membrane is implicitly defined as the region where ϕ = 0 (green line). The microtubules (bleached red lines in the background) are not tracked individually but only through the orientation field *P*, which gives the average microtubule direction at each point. The protrusive force of the microtubules is modeled by active stress in the direction of the orientation field (red arrows). The contractile myosin forces are modeled by the surface tension and bending forces (green arrows), which minimize the surface curvature and area. (**B**) Initially (at time t=0), the cell is chosen to be elliptic with microtubules oriented in the proximal-distal (x) direction. (**C**) Cell shape at steady state (time t = 5) for counteracting protrusive and contractile (surface tension) forces. The computational parameters were chosen as Fa = -1 and Ca = 0.1. For this choice of parameters, the cell approximately maintains its shape, indicating a force balance. (**D**) If the protrusive force is reduced (Fa = -10, Ca = 0.1), the cell is significantly shorter in the proximal-distal direction at a steady state. (**E**) If the surface tension is reduced instead (Fa = -1, Ca = 1), the cell is significantly elongated at a steady state. The color bar shows the value of phase-field parameter ϕ, turquoise arrows represent orientation field *P* (average microtubule orientation). The white line indicates the cell membrane (zero level set of ϕ).

As shown in Figure 4, our model recapitulates overall cell shape changes observed in vivo (Figure 2C-C’’). For the right choice of parameters, a force balance is obtained such that the cell approximately maintains its shape in steady state (Figure 4B-C). If the protrusive forces are reduced, modelling a disassembly or misorientation of the microtubules, the cell shortens along the proximal-distal axis (Figure 4D, compare lower branch of Figure 2A). On the other hand, for reduced actomyosin contractile forces (reduced effective surface tension), the cell becomes significantly elongated (Figure 4E, compare upper branch of Figure 2A).

### Local forces are shaping tissue morphology

The findings to this point argue for an interplay of microtubule protrusive and actomyosin contractile forces as a regulatory mechanism driving cell shape changes during tissue morphogenesis. To validate this observation, we systematically tested the role of microtubule and actin-based forces during wing elongation. We reasoned that changes on the cell level should translate into tissue-level changes. Strikingly, loss of MyosinII function drastically affected the tissue shape. The anterior compartment, where MyosinII function was abrogated, was significantly longer than the posterior compartment that was used as an internal wild-type control (Figure 5A,B). This is consistent with our result showing that cortical tension inversely correlates with cell elongation (Figure 1E and Figure 2B). Given that the Fat-PCP signaling pathway is required to direct microtubule-generated forces, we reasoned that in the *ft-PCP* mutant wing epithelium, where cells are shorter, the tissue would also be shorter. In agreement with our findings on the cell level, perturbation of the microtubule cytoskeleton led to a significant defect in tissue elongation (Figure 5A, bottom). Moreover, *ft-PCP* mutant tissue is similar in size and shape to the wild-type wing at the onset of cell elongation (Supp Figure 1A), further strengthening the notion that Fat-PCP-dependent patterning of forces is required for cell and tissue shape at stage I. However, to rule out unspecific effects resulting from possible interaction between the Fat-PCP pathway and MyosinII, we probed whether the global level of phosphorylated MyosinII and its polarity are preserved in *ft-PCP* mutant tissue. Despite changes in cell shape, this was the case (Figure 5C-E), revealing that the Fat-PCP signaling pathway acts upstream of the microtubule cytoskeleton in the wing epithelium but does not regulate actomyosin. Additionally, the analysis of Patronin depleted wings shows a significant reduction in wing length, confirming the specific requirements of microtubules for cell and tissue elongation (Figure 5F,G).

**Figure 5.**
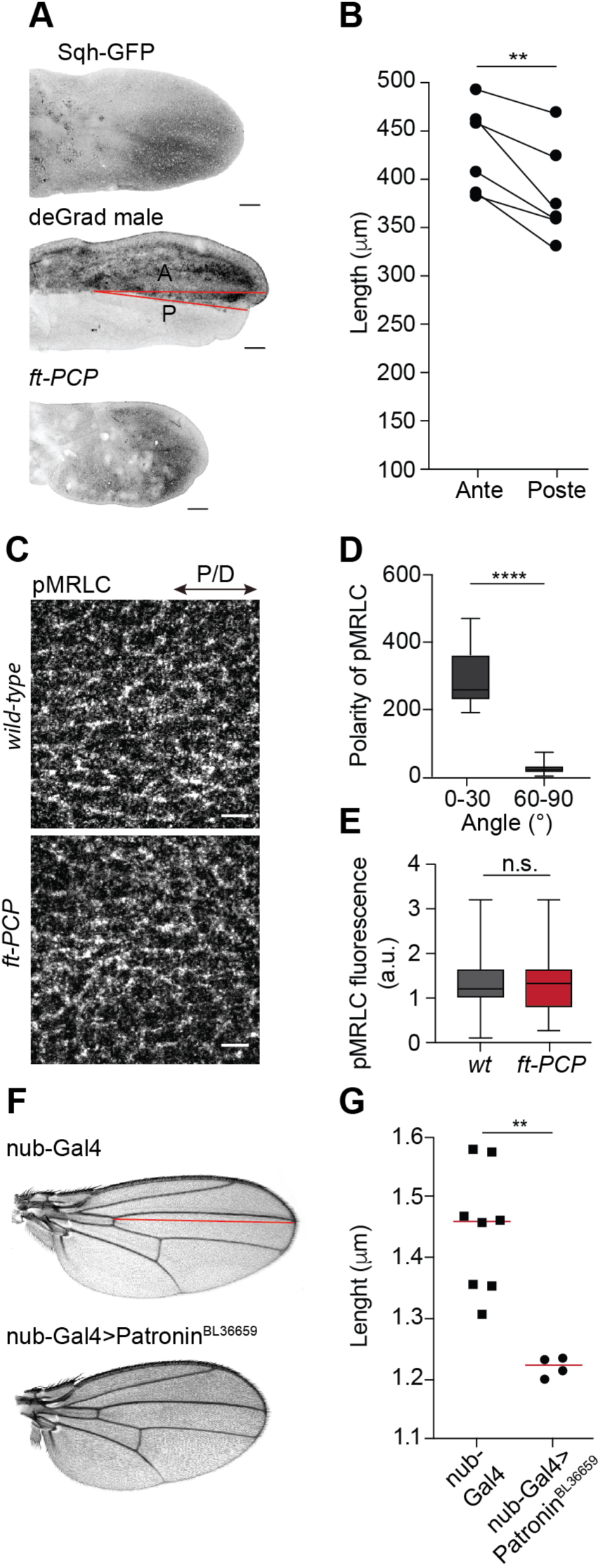
Cell shape changes affect tissue shape. (**A**) Comparison of *wild-type* (Sqh-GFP male), deGradFP male, and *ft-PCP* (*ft^l(2) fd^/ft^GRV^;tub-Gal4/UAS-Ft*Δ*ECD*Δ*N-1*) mutant wings at 18 hAPF. (**B**) Graph showing quantification of anterior and posterior wing lengths in the deGradFP male marked as indicated by red lines in (A) (Two-tailed paired t-test, ** p = 0.0063, n=6 wings and 6 pupae). Graph show scatters dot plots with the red line indicating the mean. (**C**) Representative images of 18 hAPF *wild-type (w^1118^)* wings (top) and *ft-PCP* (bottom) stained for pMRLC and Arm. (**D**) Quantification of pMRLC polarity in *ft-PCP* (*ft^l(2) fd^/ft^GRV^;tub-Gal4/UAS-Ft*Δ*ECD*Δ*N-1*) wings (Mann-Whitney test, **** p < 0.0001, n=320 cells and 3 pupae). (**E**) Quantifications of mean intensities of pMRLC in *wild-type (w^1118^)* and *ft-PCP* (*ft^l(2) fd^/ft^GRV^;tub-Gal4/UAS-Ft*Δ*ECD*Δ*N-1*) wings (Mann-Whitney test, n.s. p > 0.5360, n(*w^1118^*)=120 and n(*ft-PCP*)=94 junctions and 3-5 pupae per genotype). (**F**) Representative images of control (*nub-Gal4*) and *Patronin (nub-Gal4>Patronin^RNAi^)* depleted adult wings. (**G**) Graph showing quantification of wing lengths marked as indicated by the red line in (D) (Mann-Whitney test, ** p = 0.0040, n(*nub-Gal4*)=8 and n(*nub-Gal4>Patronin^RNAi^*)=4 wings). Graph show scatters dot plots with the red line indicating the mean. Scale bars, (A) 50 µm, and (C) 5 µm.

Taken together, our analysis of tissue morphogenesis illustrates that coordination of local forces generated by microtubules and not by actomyosin contractile forces are essential for tissue extension. We propose that gradients of Ds and Fj, constituents of the Fat-PCP signaling pathway, serve as an instructive cue at the cell and tissue level to pattern force generation.

### Initial cell shape changes are independent of extrinsic forces

Cell-extrinsic tensile and compressive forces applied to cells from external loads can also drive the remodeling of tissues. Indeed, as described above, a tissue-scale force generated by hinge contraction was reported to drive cell remodeling during phase II (Etournay et al., 2015, Ray et al., 2015, Sugimura and Ishihara, 2013, Bardet et al., 2013). To account for a possible cell-extrinsic contribution to initial cell elongation, we thus explored potential extrinsic mechanical forces in our system (i.e., phase I). To disentangle active cell forces from forces imposed extrinsically on wing cells, we severed the wing blade from the hinge at 18 hAPF (i.e., after the initial cell elongation and just before the onset of hinge contraction) and quantified the cell length (Supp Figure 6A, Video 1). We reasoned that if cell intrinsic forces drive the initial cell elongation, then cells should not shorten upon mechanical uncoupling of the blade from the hinge. Consistently, the analysis revealed that cells stayed elongated after 1 hour (Figure 6A,B). To exclude the stiff apical cuticle material as the possible mechanical influence that may cause cells to stay elongated after being mechanically isolated, we performed a TEM analysis of 16 hAPF old wings (Figure 6C). We unequivocally could see that the epithelium was already molted at this stage (Figure 6D), thus excluding a contribution of the cuticle to cell shape changes.

**Figure 6.**
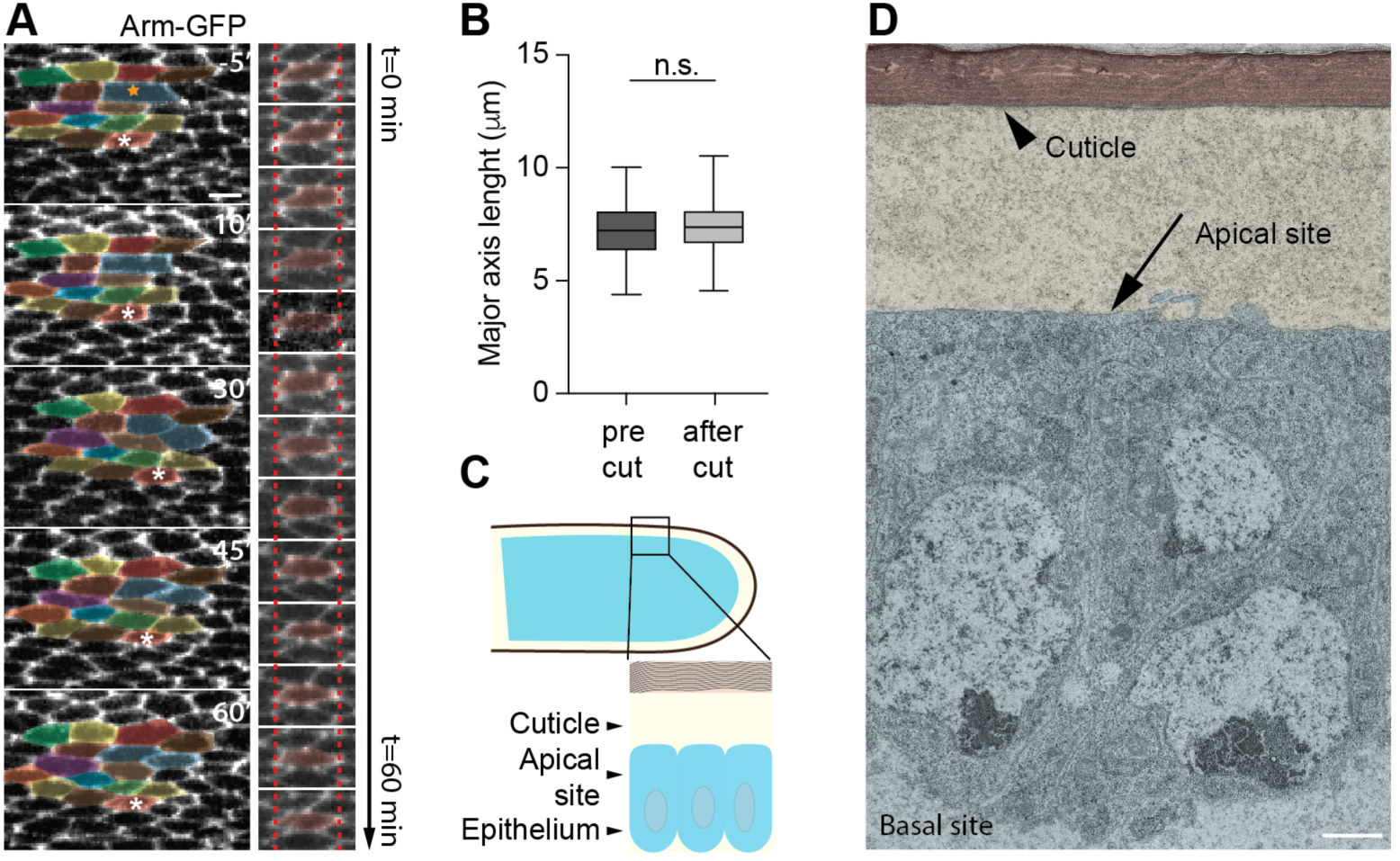
Initial wing cell elongation is independent of extrinsic forces. (**A**) Representative images of 18 hAPF wings expressing Arm-GFP 5 minutes before and after hinge cut (10, 30, 45, 60 minutes). The orange star marks the cell that will divide. (**A’**) Time-lapse images of the cell marked with the white asterisk in the (A). Note that the length of the cell does not change significantly after hinge ablation. Red dotted line marks left and right borders of the cell in the first frame. (**B**) Quantification of the cell length in 18 hAPF wings before hinge cut and after 60 minutes. (Two-tailed t-test, n.s. p > 0.1683, n(pre-cut)=136 and n(after cut)=125 cells and 3-5 pupae per genotype). (**C**) Cartoon showing a cross-section of the wing around 16 hAPF. (**D**) TEM micrograph showing a cross-section of 16 hAPF *wild-type (w^1118^)* wing. Note that the overlying cuticle (black arrowhead) is fully detached from the epithelium apical surface (black arrow). Boxes in all plots extend from the 25^th^ to 75^th^ percentiles, with a line at the median. Whiskers show min and max values. Scale bars, (A) 5 µm, and (D) 1 µm.

To further validate these findings, we analyzed *dumpy* mutants, where anchorage of the epithelium to the cuticle is perturbed. In the *dumpy* mutant, the tissue elongates normally compared to the *Ft-PCP* mutant, thus confirming that the cuticle cannot contribute to elongation until 18 hrs APF (Supp Figure 5A). Importantly, the observed mechanical autonomy of cells at this early stage is consistent with microtubule perturbation experiments, where cells fail to elongate (Figure 2B). In both the *ft-PCP* mutant and Patronin depleted wings, the hinge contracts normally (Figure 5F and Supp Figure 6B’). Hence, the initial cell elongation and consequent tissue elongation should be rescued if it depends on anisotropic stress that emerges from the hinge constriction. Since the tissue is shorter in both cases of genetic perturbation of microtubule organization (Figure 5G and Supp Figure 6B), these results provide direct experimental evidence that initial cell elongation is independent of extrinsic forces.

## Discussion

Previous studies established that microtubules contribute to cell shape changes (Singh et al., 2018, Takeda et al., 2018, Picone et al., 2010). Our observations expand this view, showing that microtubules, which are patterned along the proximal-distal axis by the Ft-PCP signaling pathway, generate protrusive forces that initiate cell elongation required for wing tissue extension. Upon polarization, actomyosin contractility further narrows these cells. Collectively, our work fills a critical gap on the interplay between local microtubule and actomyosin-generated forces during wing elongation. As depletion of MyosinII leads to the isotropic junctional tension and cell pre-stress release, it argues against the presence of additional polarized contractile force in wing cells. Consistently, downregulation of Dachs and MyosinVI, two atypical myosins previously associated with junctional remodeling (Bosveld et al., 2012, Lin et al., 2007), does not affect wing shape (Supp Figure 5B).

We further find that the patterning of non-centrosomal microtubules is independent of mechanical or geometrical cues. Cell shape was proposed to play a critical role in aligning non-centrosomal microtubules in the fly epithelium (Gomez et al., 2016). Considering their stiffness and angular dependency on catastrophe, growing microtubules could self-organize into a network controlled by the elongated cell shape (Picone et al., 2010, Mirabet et al., 2018). Given the changes in cell shape observed upon myosin inhibition, it is thus plausible to envision that actomyosin contraction acts upstream of microtubules to initiate cell elongation. However, our data revealed that despite cells becoming rounder, the alignment of microtubules along the proximal-distal axis in MyosinII depleted cells is not perturbed (Figure 2C’,E). Hence, our data support the notion that cell elongation is not the cause but the consequence of polarized alignment of microtubules via the Fat-PCP signaling pathway. Indeed, failure to pattern apical microtubule cytoskeleton upon loss of the microtubule minus-end binding protein Patronin, which is required for the correct organization of non-centrosomal microtubules in wing epithelial cell, results in shorter cells and tissue.

In summary, our results refine the current view on wing epithelium development, showing that a combination of local opposing forces drives cell elongation. We consider the identified mechanism to be complemented by additional regulatory circuits that jointly control cell and tissue shape changes during wing development. Notably this model is consistent with published data showing that extrinsic mechanical forces act during the late phase of wing reshaping starting after 18 hAPF, when the extracellular matrix protein Dumpy becomes patterned at the wing margin (Ray et al., 2015, Etournay et al., 2015). Most importantly, our findings also have implications for planar cell polarity. Ft-PCP signaling pathway, which coordinates planar polarization of cells within the tissue plane during various morphogenetic processes in invertebrates and vertebrates, is highly conserved. Considering that aberrant PCP signaling yields a failure of tissue elongation, which leads to many developmental anomalies such as body truncation and neural tube defects, we propose that microtubules and actin cytoskeleton play an important role in shaping cells during development and homeostasis. In the future, it will thus be critical to elucidate the mechanical interplay of microtubules and actomyosin and its dependency on PCP signaling in other biological contexts.

## Materials and Methods

### Drosophila Melanogaster

The following mutant and transgenic fly strains were used in this study: *w^1118^ (*BDSC 3605), *arm-Arm-GFP (*BDSC 3605), *sqh^AX3^; sqh-Sqh-GFP (*BDSC 57144), *ciGal4, tubGal4 (*BDSC 5138), *tubGal80^ts^ (*BDSC 7019), *nubGal4 (*BDSC 86108), *d^1^(*BDSC 270), *d^GC13^ (*BDSC 6389), *dp^ov1^ (*BDSC 276), *ft^GRV^ (*BDSC 1894), *ft^l(2)fd^* (BDSC 1894), *Patronin* RNAi (BDSC 36659) stocks from BDSC, *MyosinVI* RNAi stock from VDRC (VDRC 37534), *UAS-NSlmb-vhhGFP4 (UAS-deGradFP* construct) from D. Brunner, *UAS-GFP-DN-Zip* from D. Kiehart, *UAS-FtΔECDΔN1* from S. Blair. All fly stocks were raised at 25°C (unless otherwise mentioned) and grown on standard cornmeal-agar medium. The stocks are listed in Flybase (www.flybase.org).

### deGradFP expression and standardization of dual temperature regime

Continuous expression of deGradFP at 25°C was lethal for the embryos and did not allow the growth until the desired stages of development. Therefore, in order to disrupt MyosinII contractility during the desired stages of pupal wing development, deGradFP was required to be expressed in a temporally controlled way using *ciGal4* driver and tubGal80^ts^ system at a combination of 18°C and 29°C. The different time regimes of dual temperature were established in order to get the desired developmental stages where the role of MyosinII contractility was analyzed. The different dual temperature regimes, their equivalent developmental stages and the assays done using those regimes are summarized in Table 1 (Supplemental information).

### Adult wing preparation

Wings from the adult flies were dissected and mounted on a glass slide in Canada balsam (Sigma-Aldrich) mixed with a tiny drop of absolute ethanol.

### Immunohistochemistry

White pre-pupae were collected at 0 hAPF from desired stocks and crosses growing either at 18°C (for all deGradFP sets of experiments) or 25°C (for all sets of experiments other than deGradFP). The pre-pupae collected at 0 hAPF were then either shifted to 29°C (for all tubGal80^ts^ set of experiments) or grown at 25°C (for all set of experiments other than tubGal80^ts^) until the final desired developmental stages were reached. The pupae were then fixed for 6-8 hours in 4 % PFA (with 0.1% Triton X-100) at room temperature. Wings were dissected and washed thoroughly with PBT (PBS + 0.1% Triton X-100) 2-3 times. Immunostaining was performed using standard protocols (Matis et al., 2014) with minor modifications. Primary antibody (or antibody cocktail with appropriate antibody dilution) was first diluted in PBTB (PBS + 0.1% Triton X-100 + 2% of BSA) and then added to the wings. The wings were then incubated overnight (∼16 hours) at 4°C on a rotor. After three washes using PBT secondary antibody diluted in PBTB was added to the wings. Wings were incubated for 2 hours at room temperature and washed three times in PBT. Finally, wings were mounted on glass slides in Vectashield with or without DAPI (Vector Laboratories) and covered with coverslips.

For immunofluorescence staining with pMRLC, pupae were fixed for 1 hour in 18% PFA (with 0.1% Triton X-100) at room temperature. Wings were dissected and washed in PBT (PBS, 0.3% Triton X-100) for 3 times (10min each) followed by blocking for 1 hour with PBTB (PBS, 0.3% Triton X-100, 0.5% BSA) and incubated overnight in primary antibody cocktail (in PBTB) at 4°C. Wings were washed 4 times (10 min each) using PBT. Wings were then incubated in a fluorescently conjugated secondary antibody cocktail (in PBTB, for 2 hours, at room temperature) and were washed 3 times (20 min each) with PBT before mounting on glass slides in Vectashield with DAPI.

The following primary antibodies were used: rabbit anti-α-tubulin (1:200; ab18251, Abcam); mouse anti-armadillo (1:100; N2 7A1, DSHB); mouse anti-flamingo (1:100; DSHB); and rabbit anti-pMRLC (1:50; 3671S, Cell Signaling Technology). Rhodamine phalloidin dye (1:100; Invitrogen) was used for visualizing pre-hair orientation. Fluorophore-conjugated secondary antibodies (Invitrogen) were used at 1:200 dilution.

### Imaging of pupal wing

Fixed and stained wings were imaged using an upright LSM 710 Confocal microscope (Carl Zeiss). The images were acquired using Zen software (Carl Zeiss, version 6.0, 2010) at different high magnification objectives for different experiments depending on the resolution required. In general, 5x (0.16 EC Plan-Neofluar, Carl Zeiss) was used for all low magnification images to visualize whole wing morphology and determine the landmarks for precise identification of developmental stages. For high magnification images, 40x (1.3 Oil C Plan-Apochromat, Carl Zeiss), 63x (1.4 Oil Plan-Apochromat, Carl Zeiss) and 100x (1.46 Oil α-Plan-Apochromat, Carl Zeiss) objectives were used. All high magnification images were acquired at a step size of 0.3 µm (unless otherwise mentioned).

### Imaging of adult wing

Adult wings were imaged using Imager.M1 microscope (Carl Zeiss) equipped with CoolSNAP ES2 camera (Photometrics) using 10x objective (0.3 EC Plan-Neofluar, Carl Zeiss).

### Transmission Electron Microscopy (TEM)

For TEM analysis, pupal wings of the appropriate age were fixed overnight at RT in a mixture of 2,5% glutaraldehyde in phosphate buffer (pH 7.3) and were further processed as described previously (Singh et al., 2018).

### Laser ablation

Pupae of desired genotypes and developmental stage corresponding to 18 hAPF were collected and the cuticle was gently removed to get the pupae out of their pupal cases. The pupae were placed laterally on a coverslip smeared with a very thin layer of glue placed on top of a glass slide with spacers. The pupae expressed Arm-GFP or Sqh-GFP to visualize cell junctions and actomyosin cables for ablation, respectively. A single-pulse of 355 nm laser (DPSL-355/14, Rapp OptoElectronics) at 2% laser power was used across a 10-pixel (0.64 µm) line perpendicular either to the center of cell junctions (aligned along proximal-distal and anterior-posterior axes) or to the center of Myosin cable for ablation. Time-lapse 2D images with a frame rate of 100 ms were acquired using a 100x objective (1.46 NA Oil α-Plan-Apochromat, Carl Zeiss) mounted on an upright AxioImager.M2 microscope (Carl Zeiss) equipped with CSU10B spinning disk (Yokogawa) and an sCMOS ORCA Flash 4.0LT system (Hamamatsu). Images were acquired using VisiView software (Visitron Systems GmbH) from at least 1 minute prior to ablation and up to 4 to 5 minutes post ablation to visualize the movement of cell junctions, vertices and MyosinII cables before and after nano-ablation.

For laser-induced hinge ablation, a single-pulse of 355 nm laser (DPSL-355/14, Rapp OptoElectronics) at 2% laser power was used to ablate the hinge of one of the wings (roughly near the hinge-blade border) using a line ROI along its entire width such that the cells in the blade were mechanically uncoupled to the hinge. Time-lapse images at a time interval of 5 min between the frames were acquired using a 40x (1.3 NA Oil C Plan-Apochromat, Carl Zeiss) objective. Images were acquired from at least 5-10 min prior to ablation till the end of hexagonal packing corresponding to 26 hAPF in order to visualize cell elongation (at an early stage) within minutes of hinge ablation and hexagonal packing of the cells (at a late stage) upon hinge ablation.

### Image Processing

Images were processed and analyzed using Fiji/ImageJ software (NIH, version 2.0.0, 2015). Images of pupal and adult wings acquired in parts were stitched into whole wings using the Fiji plugin “pairwise stitching”. For all the analyses related to apical cell shape, microtubule orientation, pMRLC signal intensity and apical area, only the apical slices at the level of adherens junctions were z projected. The z-slices at the level of adherens junctions in different images were determined by signals from Arm-GFP, anti-Armadillo antibody, anti-α-Tubulin antibody, anti-pMRLC antibody and Sqh-GFP. All the measurements were done in Fiji using the “Analyze” feature. Brightness and contrast were adjusted within the linear range wherever needed. If used, the median filter of 0.5 or 1.0 was applied to all the images in a given set of analyses. The whole wing images were cropped in Fiji to represent appropriate ROIs wherever needed. Appropriate and desired fluorophore channels were merged or split from multi-channel images for representation as required. Images were also converted into greyscale wherever needed for representation.

### Quantitative analyses

Images processed in Fiji were used for quantitative analyses. Measurements were done in Fiji from the ROIs drawn manually for various parameters. All the raw measurements from Fiji were then summarized and further computed. The different ROIs taken from wings for all the measurements are marked in respective figure panels in the low magnification whole wing images. The total number of ‘n’ and ‘N’ analyzed for the measurements in different experiments are mentioned in the associated figure legends.

### Quantification of cell length

ROIs were drawn manually by tracing the apical cell outlines marked by Arm-GFP or anti-Armadillo antibody signals. “Feret’s diameter,” a Fiji function, was used to measure the length of cells.

### Quantification of wing length

ROIs were drawn manually between the anterior crossvein (ACV) and the end of the longitudinal L3 vein, as shown in Figure 3D. The length was measured using Fiji function “Analyze >> Area”.

### Quantification of cell area

ROIs were drawn manually by tracing the apical cell outlines marked by Arm-GFP or anti-Armadillo antibody signals. The area was measured in square microns (µm^2^) using Fiji function “Analyze >> Area”.

### Quantification of pupal vein area

The images of the whole pupal wing were z-projected at the level of maximum cross-sectional vein area. ROIs were drawn manually by tracing the edges around the vein L2. The area was measured in square microns (µm^2^) using the Fiji function “Analyze >> Area”.

### Quantification of displacement and recoil velocity of junctions upon laser ablation

The movement of vertices associated with the ablated junctions was tracked post-ablation manually for 2 seconds at an interval of 200 milliseconds. The displacement was calculated by the distance between the two vertices at every 200 milliseconds over the period of 2 seconds. This was represented as displacement-time graph with mean and error values for total of 10 measurements at every 200 milliseconds. Recoil velocity was calculated for the initial 200 milliseconds (in the linear range) by dividing initial displacement over the first 200 milliseconds. This initial recoil velocity was used as a proxy measurement for junction tension.

### Quantification of percentage angular distribution of microtubules

Appropriate apical slices were z projected. All the wings were aligned along their proximal-distal axis. Angular or polarized distribution of microtubules was measured by the Fiji plugin “Orientation J”. Microtubule orientation from different wings of same genotypes were pooled together and binned into 3 categories of angular distribution (0-30°, 30-60° and 60-90°) with the proximal-distal axis. The mean population of microtubules oriented across the three categories were compared.

### Quantification of pMRLC polarity and intensity along the junctions

Appropriate apical slices were z projected. All the wings were aligned along their proximal-distal axis. Polarization of pMRLC was measured by the Fiji plugin “Orientation J”. pMRLC orientations were pooled together and binned into 3 categories of angular distribution (0-30°, 30-60° and 60-90°) with the proximal-distal axis. The mean population of binned categories were compared. The quantification of the junctional pMRLC intensity was done by measuring the mean intensity of a 3 pixel-thick line (corresponds to 400 nm-wide stripe) using the Fiji linear ROI function along the junctions that were visualized by Arm staining. The background signal was subtracted from each of the intensity signals. Then the intensity values along each junction were normalized in respect with the average intensity signal of the control junctions in the same image (junctions on the posterior site of the wing).

### Quantification of cell shape (circularity)

ROIs were drawn manually by tracing the apical cell outlines using Arm-GFP or anti-Armadillo antibody signals. Cell circularity was measured using the Fiji function “Shape descriptors”. Cell circularity is a measure of how close a cell is to being a perfect circle (circularity = 1). Mean cell circularity for many cells was computed and compared across different genotypes.

The formula for circularity is as follows:

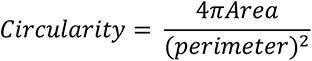

### Computational model of actomyosin-microtubule force balance

The model was originally used to model cell motility by actomyosin contractile stress (Marth et al., 2015). Here, we adopt the approach to model active forces generated by the microtubules (Figure 3A). The cell is modeled as an active polar gel surrounded by a membrane that separates it from the surrounding extracellular fluid. The model uses a diffuse interface description of the cell, i.e., the cell is modeled by the phase-field parameter ϕ that takes on the value 1 inside the cell and -1 outside with a smooth transition in the interface region. The cell membrane is implicitly defined as the region where ϕ = 0. The average orientation of the microtubules is tracked by the vector-valued orientation field *P*. Note that *P* only represents the average direction of microtubules, not a microtubule density or the strength of the local pushing force. Hence, the model (weakly) enforces unit length |*P*| = 1 for the orientation field vectors inside the cell and ensures that the orientation field vanishes (|*P*| = 0) outside the cell, with a smooth transition in the interface region. For the fluid, we track the velocity *u* and the pressure *p* using the Stokes equations. For simplicity, we assume equal density for the cytoplasm and the extracellular fluid.

The model equations are obtained by assuming that the system evolves according to a gradient descent of the free energy

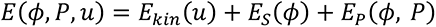

which is composed of the kinetic energy of the fluid, the membrane energy *E_S_* and the energy of the microtubule network *E_P_*. The membrane energy consists of Helfrich-type bending energy (Helfrich, 1973) and the surface energy

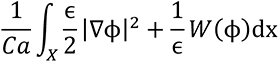

corresponding to a classic Cahn-Hilliard model (Cahn and Hilliard, 1958, Cahn and Hilliard, 1959). Here, ɛ is the phase-field parameter describing the thickness of the interface region and *W* is a double-well potential with minima at -1 and 1. The capillary number *Ca* regulates the strength of the surface tension (higher values of *Ca* model lower surface tension). The bending resistance is controlled by the parameter *Be*.

The cell (phase field) and the microtubules are advected with the fluid flow and appear as additional stress terms in the Stokes equation. In particular, the protrusive force of the microtubules enters via the active stress term

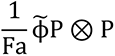

where 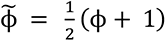 is an indicator function for the cell interior and the (negative) active force number *Fa* controls the strength of the protrusive force (higher absolute value of *Fa* means lower protrusive force). In our simulations, we keep *Be* fixed and only vary *Ca* and *Fa*, i.e., we assume a fixed bending resistance and model the actomyosin forces through the surface tension.

The resulting coupled non-linear system of partial differential equations (PDEs) is solved using our discretization module dune-gdt (https://zivgitlab.uni-muenster.de/ag-ohlberger/dune-community/dune-xt/-/blob/master/README.md) based on the software framework DUNE (Bastian et al., 2021). For a detailed description of the model and the numerical approach, see (Leibner et al., 2021).

### Statistical analyses

All the datasets presented in this work were first tested for normal distribution using D’Agostino & Pearson test or Shapiro-Wilk normality tests. Statistical significance was determined using two-tailed Unpaired/Student’s t-test or ordinary one-way ANOVA (with Tukey’s multiple comparisons) when the data were distributed normally for two or more than two groups, respectively. Two-tailed Mann-Whitney U-test or Kruskal-Wallis test (with Dunn’s multiple comparisons) were performed when datasets were not distributed normally. All the graphs with scatter dot plots show mean values with red lines. Boxes in all box plots extend from the 25^th^ to 75^th^ percentiles, with a line at the median. Whiskers show min and max values. All the bar graphs indicate mean ± sd. The significance levels in all the graphs are as follows; n.s. (non-significant), * (*p* ≤ 0.05), ** (*p* ≤ 0.01), *** (*p* ≤ 0.001) and **** (*p* ≤ 0.0001). The statistical tests were performed using Prism7 (version 7.0d for Mac OS X, GraphPad Software). All experiments presented in the manuscript were repeated at least in three independent experiments/biological replicates. The experiments were not randomized, and the sample size was not predetermined.

### Data and code availability

All data are available in the main text or the supplementary materials. Code generated for this study is available from the corresponding author without restriction.

## Acknowledgments

We thank Damian Brunner, Daniel P. Kiehart, Seth S. Blair, the Bloomington Stock Center and the Developmental Studies Hybridoma Bank, for providing fly stocks and antibodies; Matis lab members for critical reading of the manuscript. This work was supported by the Cells-in-Motion Cluster of Excellence EXC-1003 (FF-2015-07) and the Deutsche Forschungsgemeinschaft (DFG, SPP-1782, MA 6726/3-1) to M.M., by the DFG under Germany’s Excellence Strategy EXC 1003 FF-2015-07 and EXC 2044 – 390685587, Mathematics Münster: Dynamics–Geometry–Structure to T.L. and M.O.

## Author Contribution

A.S. designed, performed experiments and analyzed the data, and wrote the manuscript. S.T. performed the immunostaining experiments. A.R., H.N. and J.K. prepared and imaged TEM samples. T.L. and M.O. developed the model and carried out the simulation. M.M. supervised the research and wrote the manuscript with feedback from all authors.

**Video 1. Hinge ablation experiment.**

Live imaging of hinge region of 18 hAPF pupal wing expressing Arm-GFP before and after ablation. Scale bar, 25 μm.

---

**Supplemental Figure 1.**
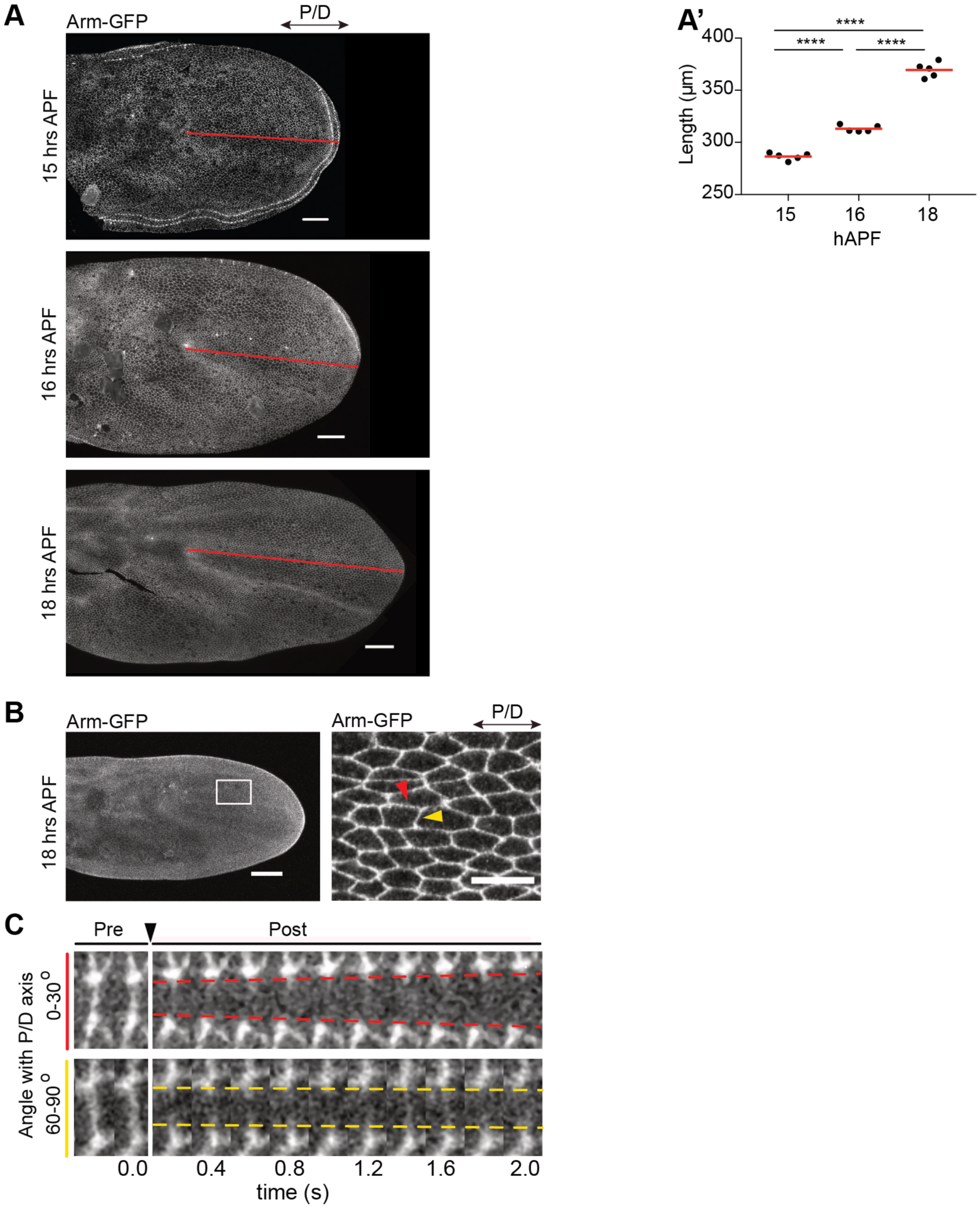
Wing tissue elongates between 14 and 18 hAPF. (A) During pupal wing morphogenesis, the epithelium undergoes cell shape changes (between 14 to 18 hAPF) that are critical for the elongation of the wing. (A’) Graph showing quantification of wing lengths marked as indicated by red lines in (A) (Ordinary one-way ANOVA, from left to right: **** p < 0.0001, **** p < 0.0001, **** p < 0.0001, n=5 pupae per genotype). Scale bars, 25 µm. (B) 18 hAPF pupal wing grown at 25°C expressing Arm-GFP. The right panel shows the boxed region from left panel. Red arrowhead marks cell junction oriented along proximal-distal and yellow arrowhead marks anterior-posterior oriented junction. (C) Kymograph showing displacement of vertices upon laser ablation of cell junctions along proximal-distal (top, red dashed lines) and anterior-posterior (bottom, yellow dashed lines) axes. Black arrowhead points to the time of laser ablation. Notice that the displacement of vertices for junction along the proximal-distal axis is larger compared to that along the anterior-posterior axis in the same period of time.

**Supplemental Figure 2.**
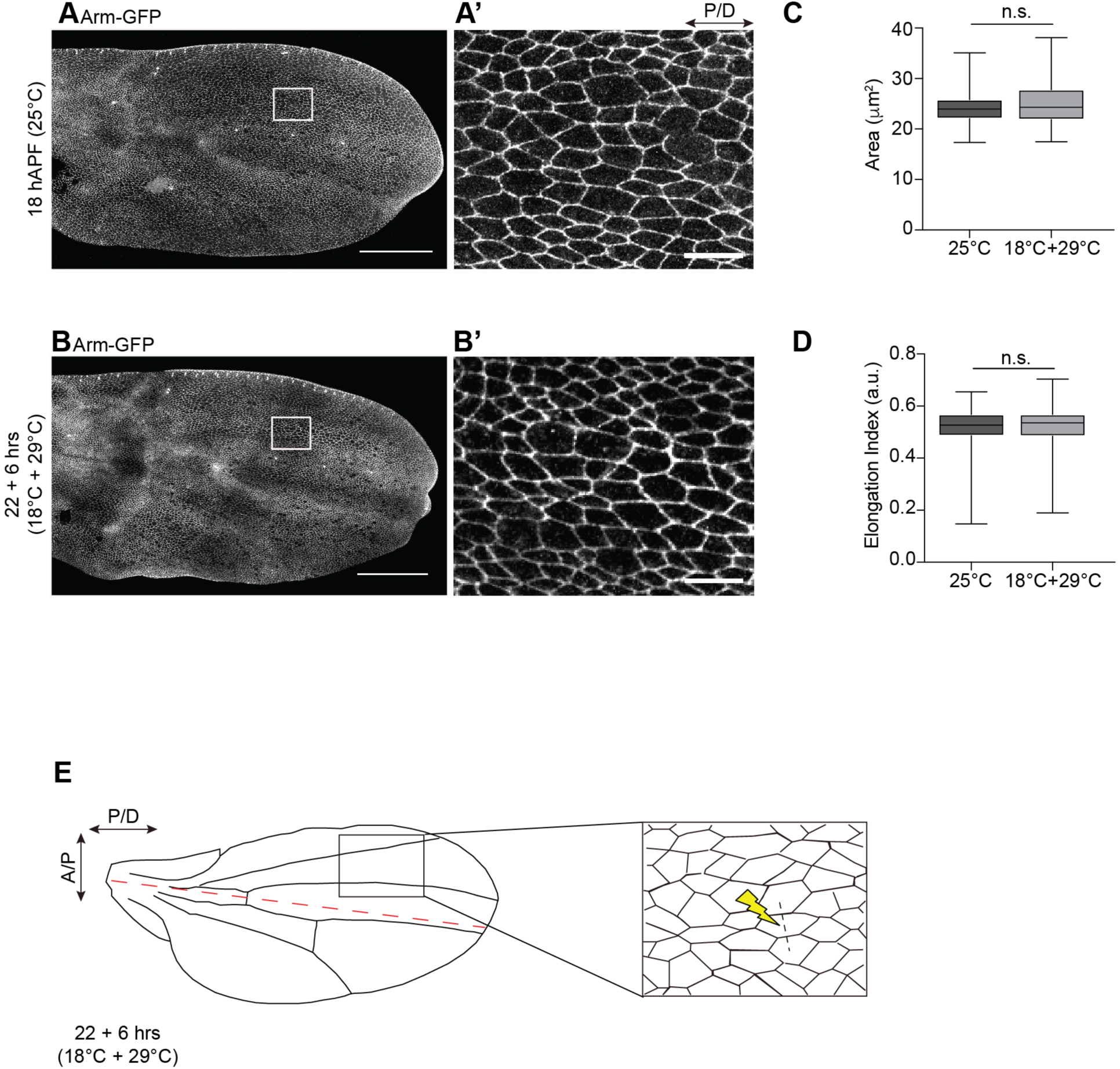
Standardization of growth conditions for the developmental stage (∼18 hAPF) at dual temperature regime. (A) 18 hAPF pupal wing grown at 25°C expressing Arm-GFP. (A’) shows the boxed region from (A). (B) Arm-GFP wing grown at a combination of 18°C and 29°C dual temperature regime using the tub-Gal80^ts^ system to get developmental stage equivalent to 18 hAPF at 25°C. (B’) shows the boxed region from (B). (C and D) Quantification of apical cell area (C) and elongation index (D) for Arm-GFP expressing flies grown at two different regimes. (C and D: Two-tailed, Mann-Whitney U-tests, n.s. (for C) p = 0.2374, n.s. (for D) p = 0.8631, n=150 cells and 3 pupae each for both the regimes). Boxes in all plots extend from the 25^th^ to 75^th^ percentiles, with a line at the median. Whiskers show min and max values. Images shown in (A-B’) are representative of n=3 wings and N=3 independent experiments. Scale bars, (A and B) 100 µm, (A’ and B’) 10 µm. (E) Cartoon showing pupal wing grown at a combination of 18°C and 29°C dual temperature regime using the tub-Gal80^ts^ system to get developmental stage equivalent to 18 hAPF at 25°C (top) and inset of cells from anterior compartment of the corresponding wing (bottom). The red dashed line (top) indicates the anterior-posterior wing boundary. Yellow thunder and black dashed line (bottom) indicate a scheme for laser ablation.

**Supplemental Figure 3.**
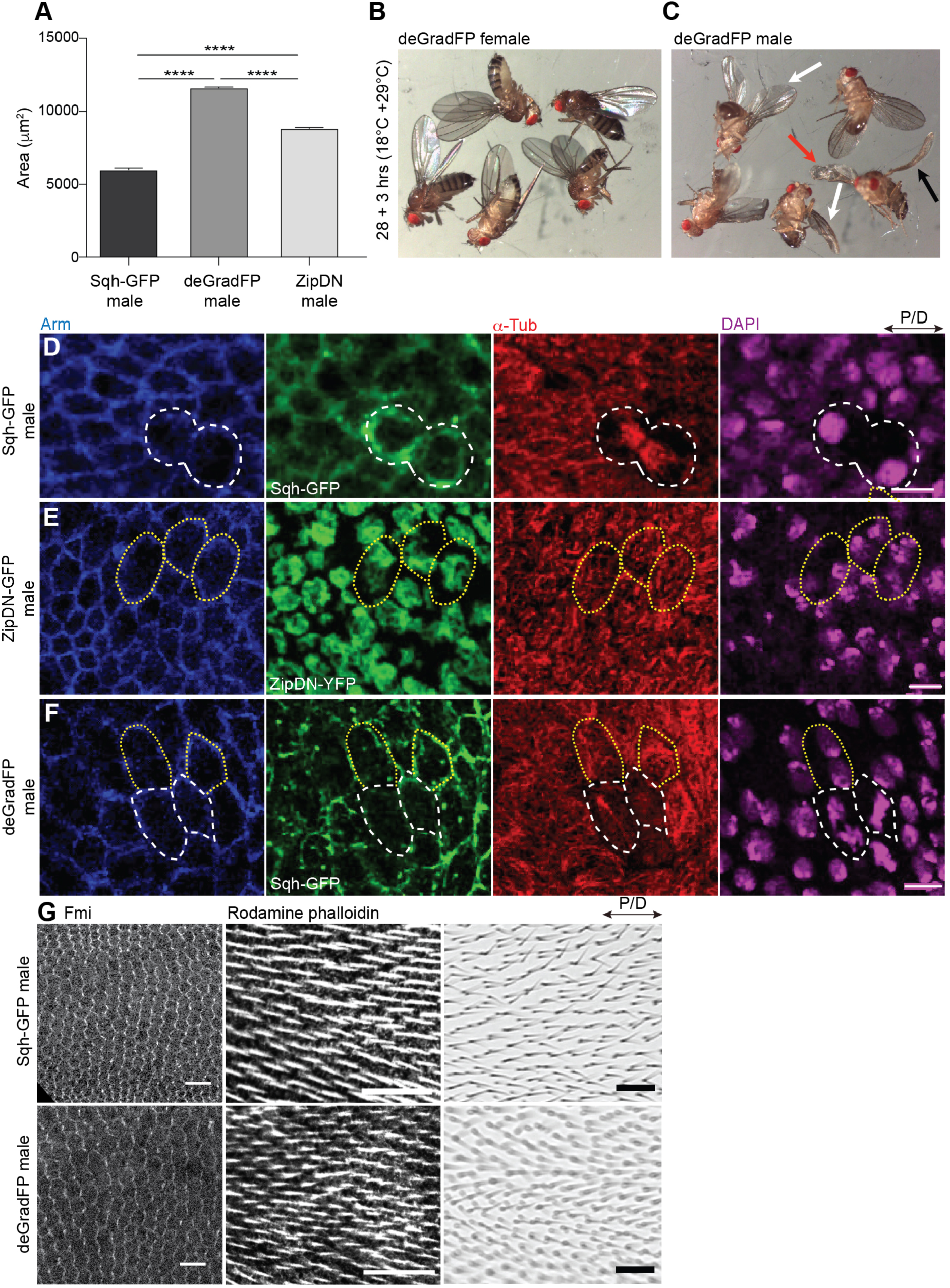
deGradFP-mediated knockdown of Sqh-GFP produces known phenotypes consistent with loss of MyosinII function in *Drosophila* pupal wing (i.e., vein expansion, blisters and aberrant cell division). (A) Quantification of L2 vein area for control wings and MyosinII depleted wings (deGradFP male and ZipDN-GFP male) (Ordinary one-way ANOVA, **** p < 0.0001, n=3 veins and 3 pupae per genotype). The graph shows bars indicating mean and sd. (B and C) Images showing deGradFP female (B) and deGradFP male (C) adult flies grown at a combination of 18°C and 29°C temperature regime using the tub-Gal80^ts^ system. Although adult wing hair orientation appears unaffected upon deGradFP expression (see panel G), note that some other defects are visible. With as low as 3 hrs of deGradFP expression, deGradFP male wings (C) show defects such as blisters (red arrow), curled up (black arrows) and droopy wings (white arrows). In contrast, deGradFP female (B) adult flies raised under the same conditions do not show any obvious wing phenotype(s) and can fly normally. Images shown in (B,C) are representative of 10 adult flies and 3 independent experiments. (D-F) Images showing cell division event(s) in control flies (Sqh-GFP male, D), ZipDN-GFP male (E) and deGradFP male (F). Cells express Sqh-GFP (D,F) and ZipDN-GFP (E) and are stained with anti-Arm antibody to visualize cell outlines apically, anti-α-Tub antibody to visualize microtubules and DAPI to visualize nuclei in the cells. Cells in Sqh-GFP male (D) have two nuclei only when the cells undergo division, as evident by the presence of spindle microtubules, MyosinII enrichment at cleavage sites and rounded-up cell morphology (all indicated by broken white lines within the corresponding panels). However, cells in ZipDN-GFP male (E) and deGradFP male (F) show two nuclei (yellow dotted lines) even when the cells are not undergoing division as evident by the absence of spindle microtubules and presence of apical planar microtubules (yellow dotted lines) and non-rounded or elongated cell morphology (yellow dotted lines) when compared to cells undergoing division where spindle microtubules (white broken lines) and nuclear division (white broken lines) are observed. Images shown in D-F are representative of n=4 pupae and N=3 independent experiments. (G) Representative images of Sqh-GFP male and deGradFP male wigs equivalent to 24 hAPF (left panels), 30 hAPF (middle panels) at 25°C and in adult animals showing that wing PCP is independent of Myosinll activity. Wings were stained with anti-Flamingo antibody to visualize PCP organization and with Rhodamine phalloidin to visualize pre-hair orientation. Images shown are representative of n=6 wings and N=3 independent experiments. Scale bars, (D-F) 5 µm, (G, left) 10 µm, (G, middle, right) 20 µm.

**Supplemental Figure 4.**
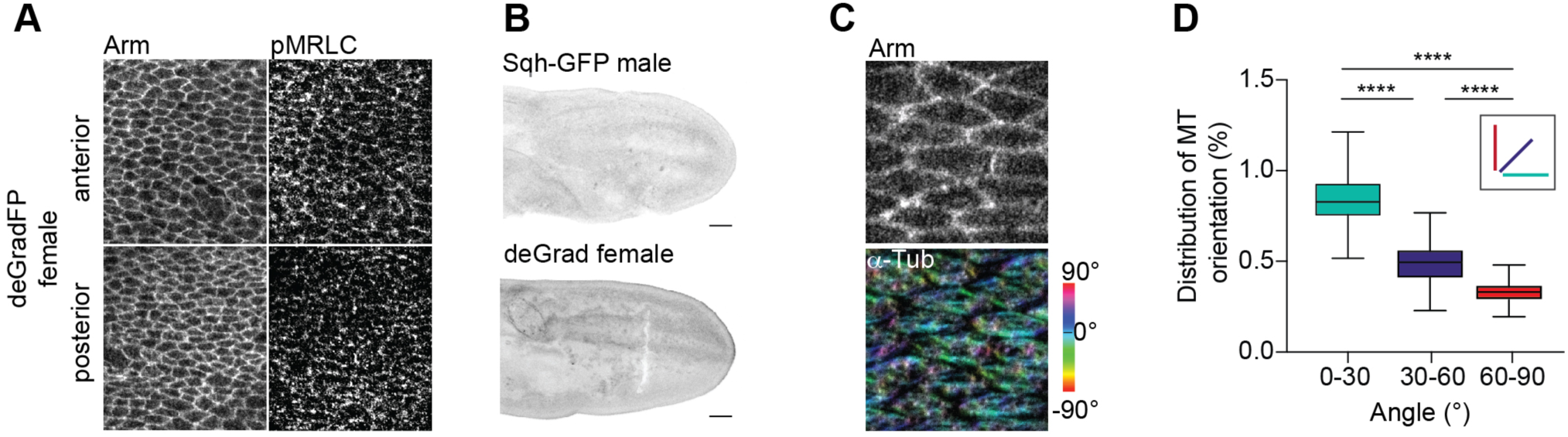
DeGradFP specifically targets Sqh-GFP and shows no artifacts. (A) Images showing anterior (top) and posterior (bellow) compartment in deGradFP female control flies (having one wild-type and one mutant copy of null mutation of *sqh^AX3^*) stained for Arm and pMRLC. In deGradFP female flies, phosphomyosin localizes to the junctions in the anterior and posterior compartments. (B) Representative images of Sqh-GFP male and deGradFP female wings at 18 hAPF show no difference. (C) Images of 18 hAPF wings of Sqh-GFP male (top) and deGradFP female (bottom) stained for Arm and α-Tub to visualize microtubules. The orientation of microtubules is color-coded using OrientationJ. The images shown are representative of 4 wings and 3 independent experiments. (D) Quantification of microtubule alignment along proximal-distal axis for deGradFP female (Kruskal-Wallis tests, from left to right: **** p < 0.0001, **** p < 0.0001, **** p < 0.0001, n=80-100 cells and 3 pupae). Boxes in the plot extend from the 25^th^ to 75^th^ percentiles, with a line at the median. Whiskers show min and max values.

**Supplemental Figure 5.**
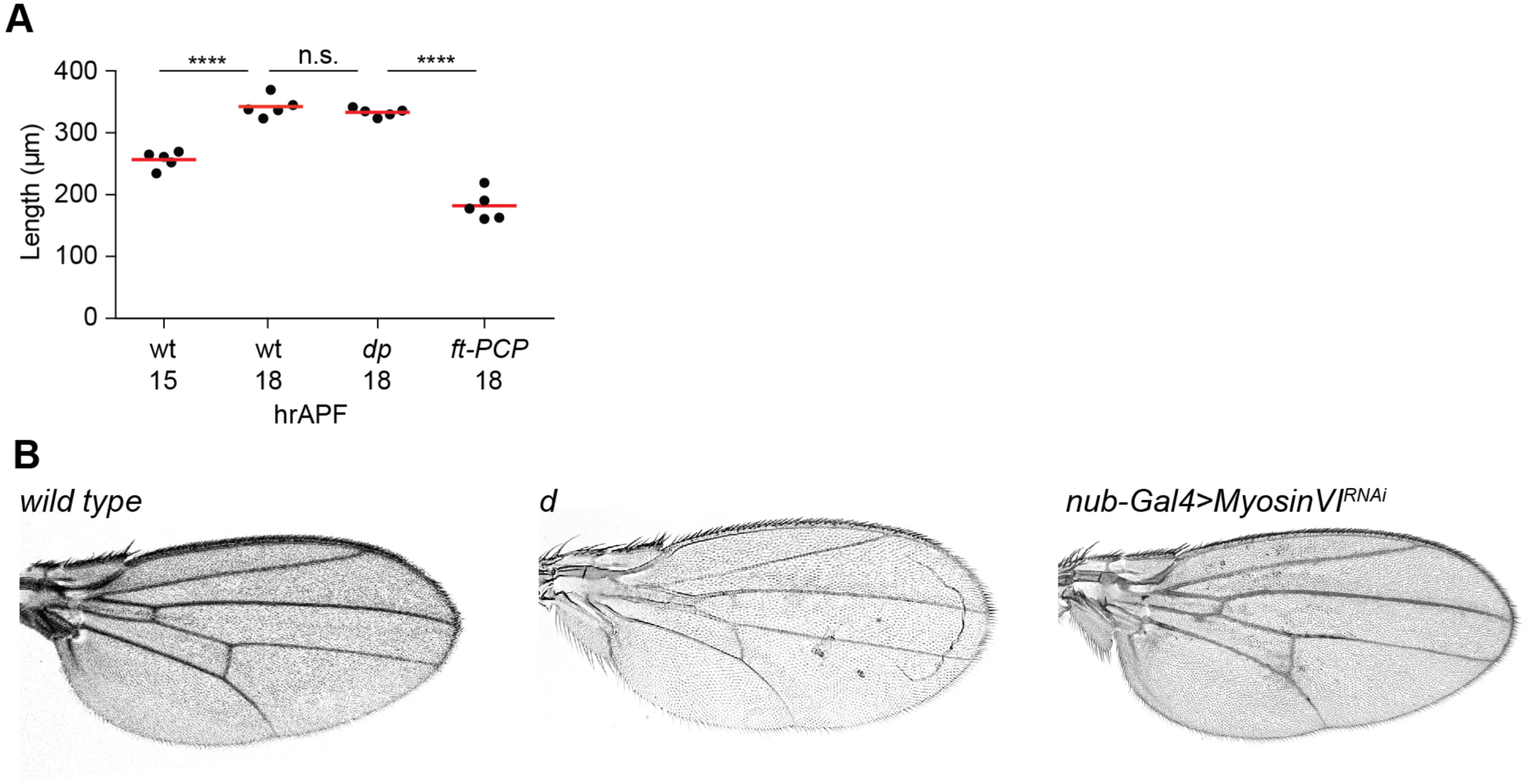
Cell shape changes affect tissue shape. (A) Graph showing quantification of 15 and 18 hAPF wild-type wing and *dumpy* and *ft-PCP* 18 hAPF mutant wing lengths marked as indicated by red lines in (Figure 1 - figure supplement 1) (Ordinary one-way ANOVA: n.s. p > 0.8127, **** p < 0.0001, n=5 pupae per genotype). Note that between 15 and 18 hAPF, wing epithelium undergoes significant extension. Likewise, the wing elongates normally in *dumpy* mutant animals, where the generation of global tissue stress is blocked. In the *ft-PCP* mutant, where cells fail to elongate, tissue is shorter. (B) Representative images of wild-type (*w^1118^*), *dachs* mutant (*d^1^/d^GC13^*) and *MyosinVI (nub-Gal4>MyosinVI^RNAi^)* depleted wings showed no defects in tissue elongation. Number of analyzed wings (*wt*)=16/16, n(*d*)= 40/40 and n(*MyosinVI*^RNAi^)=14. However, *dachs* mutant wings display abnormal vein patterning and knockdown of MyosinVI causes wing blisters.

**Supplemental Figure 6.**
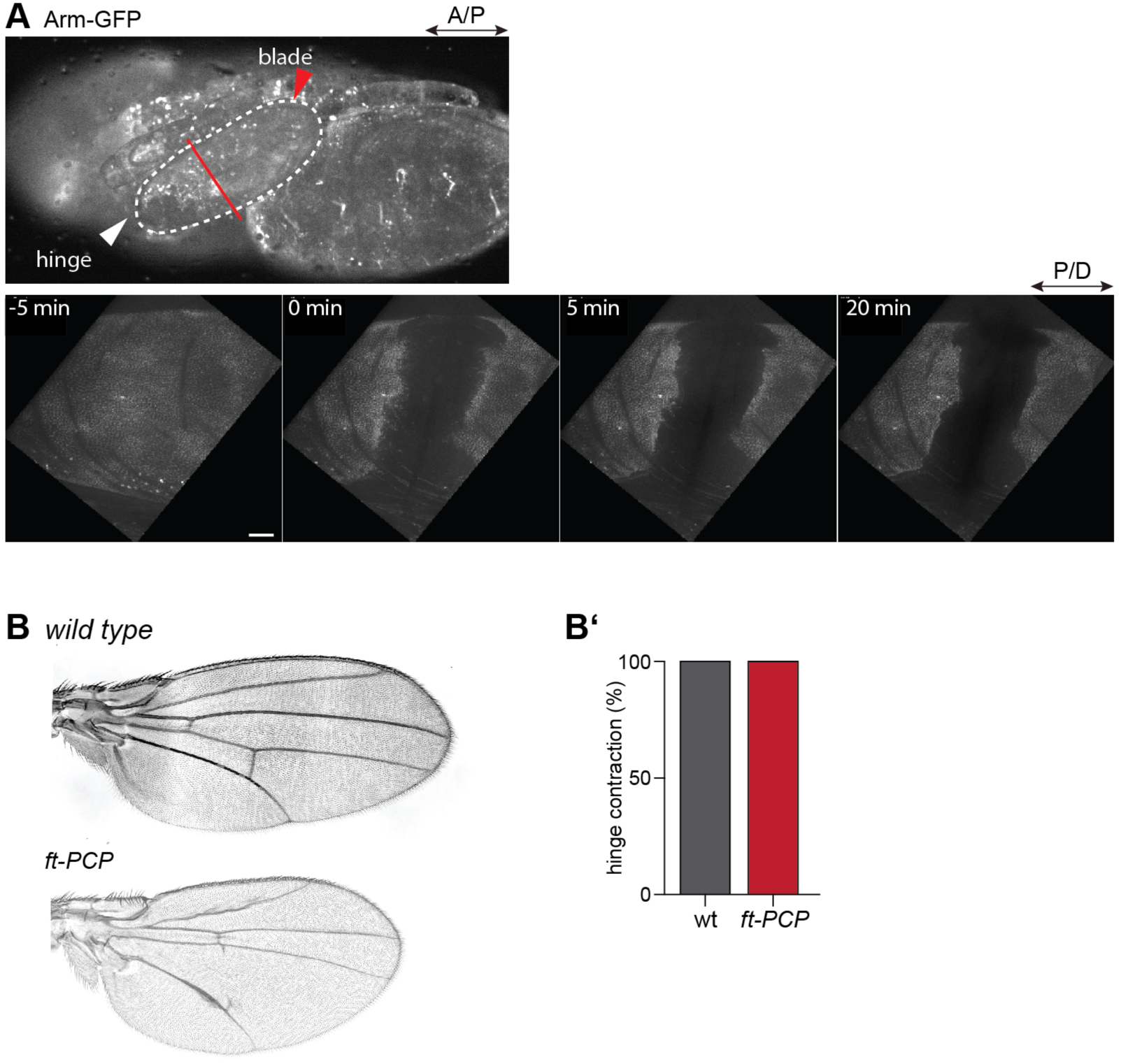
Initial wing cell elongation is independent of extrinsic forces. (A) Hinge ablation experiment (see Supplementary Video 1), showing whole pupae expressing Arm-GFP (top). The white dashed line marks the wing edge and the red line position of the laser cut. Below are time-lapse images of Arm-GFP before (5 min), during and after ablation of the wing (5 min and 20 min). (B) Representative images of wild-type and *ft-PCP* mutant wings showing normal hinge contraction. (B’) Quantification of hinge contraction in wild-type and *ft-PCP* wings. In wild-type flies, 39/39 wings display normal hinge contraction and in *ft-PCP* adults, 47/47 wings. Scale bars, (A) 25 µm.

## References

1. Aigouy, B., Farhadifar, R., Staple, D. B., Sagner, A., Roper, J. C., Julicher, F. & Eaton, S. 2010. Cell flow reorients the axis of planar polarity in the wing epithelium of Drosophila. Cell, 142, 773–86.

2. Alt, S., Ganguly, P. & Salbreux, G. 2017. Vertex models: from cell mechanics to tissue morphogenesis. Philosophical Transactions of the Royal Society B-Biological Sciences, 372.

3. Baena-Lopez, L. A., Baonza, A. & Garcia-Bellido, A. 2005. The orientation of cell divisions determines the shape of Drosophila organs. Current biology : Cb, 15, 1640–4.

4. Bardet, P. L., Guirao, B., Paoletti, C., Serman, F., Leopold, V., Bosveld, F., Goya, Y., Mirouse, V., Graner, F. & Bellaiche, Y. 2013. PTEN controls junction lengthening and stability during cell rearrangement in epithelial tissue. Dev Cell, 25, 534–46.

5. Bastian, P., Blatt, M., Dedner, A., Dreier, N. A., Engwer, C., Fritze, R., Graser, C., Gruninger, C., Kempf, D., Klofkorn, R., Ohlberger, M. & Sander, O. 2021. The DUNE framework: Basic concepts and recent developments. Computers & Mathematics with Applications, 81, 75–112.

6. Bosveld, F., Bonnet, I., Guirao, B., Tlili, S., Wang, Z., Petitalot, A., Marchand, R., Bardet, P. L., Marcq, P., Graner, F. & Bellaiche, Y. 2012. Mechanical control of morphogenesis by Fat/Dachsous/Four-jointed planar cell polarity pathway. Science, 336, 724–7.

7. Bouchet, B. P. & Akhmanova, A. 2017. Microtubules in 3D cell motility. J Cell Sci, 130, 39–50.

8. Cahn, J. W. & Hilliard, J. E. 1958. Free Energy of a Nonuniform System .1. Interfacial Free Energy. Journal of Chemical Physics, 28, 258–267.

9. Cahn, J. W. & Hilliard, J. E. 1959. Free Energy of a Nonuniform System .3. Nucleation in a 2-Component Incompressible Fluid. Journal of Chemical Physics, 31, 688–699.

10. Caussinus, E., Kanca, O. & Affolter, M. 2011. Fluorescent fusion protein knockout mediated by anti-GFP nanobody. Nat Struct Mol Biol, 19, 117–21.

11. Chen, J., Castelvecchi, G. D., Li-Villarreal, N., Raught, B., Krezel, A. M., Mcneill, H. & Solnica-Krezel, L. 2018. Atypical Cadherin Dachsous1b Interacts with Ttc28 and Aurora B to Control Microtubule Dynamics in Embryonic Cleavages. Dev Cell, 45, 376–391 e5.

12. Clarke, D. N. & Martin, A. C. 2021. Actin-based force generation and cell adhesion in tissue morphogenesis. Curr Biol, 31, R667–R680.

13. Classen, A. K., Anderson, K. I., Marois, E. & Eaton, S. 2005. Hexagonal packing of Drosophila wing epithelial cells by the planar cell polarity pathway. Developmental cell, 9, 805–17.

14. Collinet, C. & Lecuit, T. 2021. Programmed and self-organized flow of information during morphogenesis. Nat Rev Mol Cell Biol, 22, 245–265.

15. Durst, R., Sauls, K., Peal, D. S., Devlaming, A., Toomer, K., Leyne, M., Salani, M., Talkowski, M. E., Brand, H., Perrocheau, M., Simpson, C., Jett, C., Stone, M. R., Charles, F., Chiang, C., Lynch, S. N., Bouatia-Naji, N., Delling, F. N., Freed, L. A., Tribouilloy, C., LE Tourneau, T., Lemarec, H., Fernandez-Friera, L., Solis, J., Trujillano, D., Ossowski, S., Estivill, X., Dina, C., Bruneval, P., Chester, A., Schott, J. J., Irvine, K. D., Mao, Y., Wessels, A., Motiwala, T., Puceat, M., Tsukasaki, Y., Menick, D. R., Kasiganesan, H., Nie, X., Broome, A. M., Williams, K., Johnson, A., Markwald, R. R., Jeunemaitre, X., Hagege, A., Levine, R. A., Milan, D. J., Norris, R. A. & Slaugenhaupt, S. A. 2015. Mutations in DCHS1 cause mitral valve prolapse. Nature, 525, 109–13.

16. Etienne-Manneville, S. 2013. Microtubules in cell migration. Annu Rev Cell Dev Biol, 29, 471–99.

17. Etournay, R., Popovic, M., Merkel, M., Nandi, A., Blasse, C., Aigouy, B., Brandl, H., Myers, G., Salbreux, G., Julicher, F. & Eaton, S. 2015. Interplay of cell dynamics and epithelial tension during morphogenesis of the Drosophila pupal wing. Elife, 4, e07090.

18. Fan, H. F. & Li, S. F. 2015. Modeling microtubule cytoskeleton via an active liquid crystal elastomer model. Computational Materials Science, 96, 559–566.

19. Gilmour, D., Rembold, M. & Leptin, M. 2017. From morphogen to morphogenesis and back. Nature, 541, 311–320.

20. Gomez, J. M., Chumakova, L., Bulgakova, N. A. & Brown, N. H. 2016. Microtubule organization is determined by the shape of epithelial cells. Nat Commun, 7, 13172.

21. Harumoto, T., Ito, M., Shimada, Y., Kobayashi, T. J., Ueda, H. R., Lu, B. & Uemura, T. 2010. Atypical cadherins Dachsous and Fat control dynamics of noncentrosomal microtubules in planar cell polarity. Developmental cell, 19, 389–401.

22. Heisenberg, C. P. & Bellaiche, Y. 2013. Forces in tissue morphogenesis and patterning. Cell, 153, 948–62.

23. Helfrich, W. 1973. Elastic Properties of Lipid Bilayers - Theory and Possible Experiments. *Zeitschrift Fur Naturforschung C-a Journal of Biosciences*, C 28, 693–703.

24. Ikebe, M. & Hartshorne, D. J. 1985. Phosphorylation of smooth muscle myosin at two distinct sites by myosin light chain kinase. J Biol Chem, 260, 10027–31.

25. Leibner, T., Matis, M., Ohlberger, M. & Rave, S. 2021. Distributed model order reduction of a model for microtubule-based cell polarization using HAPOD. arXiv preprint arXiv:2111.00129.

26. Li-Villarreal, N., Forbes, M. M., Loza, A. J., Chen, J., Ma, T., Helde, K., Moens, C. B., Shin, J., Sawada, A., Hindes, A. E., Dubrulle, J., Schier, A. F., Longmore, G. D., Marlow, F. L. & Solnica-Krezel, L. 2015. Dachsous1b cadherin regulates actin and microtubule cytoskeleton during early zebrafish embryogenesis. Development, 142, 2704–18.

27. Lin, H. P., Chen, H. M., Wei, S. Y., Chen, L. Y., Chang, L. H., Sun, Y. J., Huang, S. Y. & Hsu, J. C. 2007. Cell adhesion molecule Echinoid associates with unconventional myosin VI/Jaguar motor to regulate cell morphology during dorsal closure in Drosophila. Dev Biol, 311, 423–33.

28. Ma, D., Yang, C. H., Mcneill, H., Simon, M. A. & Axelrod, J. D. 2003. Fidelity in planar cell polarity signalling. Nature, 421, 543–7.

29. Mao, Y., Kuta, A., Crespo-Enriquez, I., Whiting, D., Martin, T., Mulvaney, J., Irvine, K. D. & Francis-West, P. 2016. Dchs1-Fat4 regulation of polarized cell behaviours during skeletal morphogenesis. Nat Commun, 7, 11469.

30. Mao, Y., Mulvaney, J., Zakaria, S., Yu, T., Morgan, K. M., Allen, S., Basson, M. A., Francis-West, P. & Irvine, K. D. 2011. Characterization of a Dchs1 mutant mouse reveals requirements for Dchs1-Fat4 signaling during mammalian development. Development, 138, 947–57.

31. Marth, W., Praetorius, S. & Voigt, A. 2015. A mechanism for cell motility by active polar gels. J R Soc Interface, 12.

32. Matis, M. 2020. The Mechanical Role of Microtubules in Tissue Remodeling. Bioessays, 42, e1900244.

33. Matis, M., Russler-Germain, D. A., Hu, Q., Tomlin, C. J. & Axelrod, J. D. 2014. Microtubules provide directional information for core PCP function. Elife, 3, e02893.

34. Mirabet, V., Krupinski, P., Hamant, O., Meyerowitz, E. M., Jonsson, H. & Boudaoud, A. 2018. The self-organization of plant microtubules inside the cell volume yields their cortical localization, stable alignment, and sensitivity to external cues. PLoS Comput Biol, 14, e1006011.

35. Nashchekin, D., Fernandes, A. R. & St Johnston, D. 2016. Patronin/Shot Cortical Foci Assemble the Noncentrosomal Microtubule Array that Specifies the Drosophila Anterior-Posterior Axis. Dev Cell, 38, 61–72.

36. Noordstra, I., Liu, Q., Nijenhuis, W., Hua, S., Jiang, K., Baars, M., Remmelzwaal, S., Martin, M., Kapitein, L. C. & Akhmanova, A. 2016. Control of apico-basal epithelial polarity by the microtubule minus-end-binding protein CAMSAP3 and spectraplakin ACF7. J Cell Sci, 129, 4278–4288.

37. Olofsson, J., Sharp, K. A., Matis, M., Cho, B. & Axelrod, J. D. 2014. Prickle/spiny-legs isoforms control the polarity of the apical microtubule network in planar cell polarity. Development, 141, 2866–74.

38. Panzade, S. & Matis, M. 2021. The Microtubule Minus-End Binding Protein Patronin Is Required for the Epithelial Remodeling in the Drosophila Abdomen. Front Cell Dev Biol, 9, 682083.

39. Pasakarnis, L., Frei, E., Caussinus, E., Affolter, M. & Brunner, D. 2016. Amnioserosa cell constriction but not epidermal actin cable tension autonomously drives dorsal closure. Nat Cell Biol, 18, 1161–1172.

40. Penta, R., Ambrosi, D. & Shipley, R. J. 2014. Effective Governing Equations for Poroelastic Growing Media. Quarterly Journal of Mechanics and Applied Mathematics, 67, 69–91.

41. Picone, R., Ren, X., Ivanovitch, K. D., Clarke, J. D., Mckendry, R. A. & Baum, B. 2010. A polarised population of dynamic microtubules mediates homeostatic length control in animal cells. PLoS Biol, 8, e1000542.

42. Pinheiro, D. & Bellaiche, Y. 2018. Mechanical Force-Driven Adherens Junction Remodeling and Epithelial Dynamics. Dev Cell, 47, 3–19.

43. Ray, R. P., Matamoro-Vidal, A., Ribeiro, P. S., Tapon, N., Houle, D., Salazar-Ciudad, I. & Thompson, B. J. 2015. Patterned Anchorage to the Apical Extracellular Matrix Defines Tissue Shape in the Developing Appendages of Drosophila. Dev Cell, 34, 310–22.

44. Saburi, S., Hester, I., Fischer, E., Pontoglio, M., Eremina, V., Gessler, M., Quaggin, S. E., Harrison, R., Mount, R. & Mcneill, H. 2008. Loss of Fat4 disrupts PCP signaling and oriented cell division and leads to cystic kidney disease. Nature genetics, 40, 1010–5.

45. Singh, A., Saha, T., Begemann, I., Ricker, A., Nüsse, H., Thorn-Seshold, O., Klingauf, J., Galic, M. & Matis, M. 2018. Polarized Microtubule Dynamics Directs Cell Mechanics and Coordinates Forces during Epithelial Morphogenesis. Nature Cell Biology.

46. Stehbens, S. & Wittmann, T. 2012. Targeting and transport: how microtubules control focal adhesion dynamics. J Cell Biol, 198, 481–9.

47. Sugimura, K. & Ishihara, S. 2013. The mechanical anisotropy in a tissue promotes ordering in hexagonal cell packing. Development, 140, 4091–101.

48. Takeda, M., Sami, M. M. & Wang, Y. C. 2018. A homeostatic apical microtubule network shortens cells for epithelial folding via a basal polarity shift. Nat Cell Biol, 20, 36–45.

49. Toya, M., Kobayashi, S., Kawasaki, M., Shioi, G., Kaneko, M., Ishiuchi, T., Misaki, K., Meng, W. & Takeichi, M. 2016. CAMSAP3 orients the apical-to-basal polarity of microtubule arrays in epithelial cells. Proc Natl Acad Sci U S A, 113, 332–7.

50. Villano, J. L. & Katz, F. N. 1995. four-jointed is required for intermediate growth in the proximal-distal axis in Drosophila. Development, 121, 2767–77.

51. Yang, C., Axelrod, J. D. & Simon, M. A. 2002. Regulation of Frizzled by Fat-like Cadherins during Planar Polarity Signaling in the Drosophila Compound Eye. Cell, 108, 675–88.

52. Zakaria, S., Mao, Y., Kuta, A., Ferreira De Sousa, C., Gaufo, G. O., Mcneill, H., Hindges, R., Guthrie, S., Irvine, K. D. & Francis-West, P. H. 2014. Regulation of neuronal migration by Dchs1-Fat4 planar cell polarity. Curr Biol, 24, 1620–7.

53. Zeidler, M. P., Perrimon, N. & Strutt, D. I. 2000. Multiple roles for four-jointed in planar polarity and limb patterning. Developmental biology, 228, 181–96.

